# Asymmetry of cortical functional hierarchy in humans and macaques suggests phylogenetic conservation and adaptation

**DOI:** 10.1101/2021.11.03.466058

**Authors:** Bin Wan, Şeyma Bayrak, Ting Xu, H. Lina Schaare, Richard A.I. Bethlehem, Boris C. Bernhardt, Sofie L. Valk

**Affiliations:** Otto Hahn Group Cognitive Neurogenetics, Max Planck Institute for Human Cognitive and Brain Sciences, Leipzig, Germany; International Max Planck Research School on Neuroscience of Communication: Function, Structure, and Plasticity (IMPRS NeuroCom), Leipzig, Germany; Department of Cognitive Neurology, University Hospital Leipzig and Faculty of Medicine, University of Leipzig, Leipzig, Germany; Institute of Neuroscience and Medicine (INM-7: Brain and Behavior), Research Centre Jülich, Jülich, Germany; Institute of Systems Neuroscience, Heinrich Heine University Düsseldorf, Düsseldorf, Germany; Center for the Developing Brain, Child Mind Institute, New York, NY, USA; Department of Psychiatry, University of Cambridge, Cambridge, UK; McConnell Brain Imaging Centre, Montréal Neurological Institute and Hospital, McGill University, Montréal, QC, Canada

**Author notes:** Correspondence to Bin Wan and Sofie L. Valk, Otto Hahn Group Cognitive Neurogenetics, Max Planck Institute for Human Cognitive and Brain Sciences, Leipzig, Germany; Institute of Neuroscience and Medicine (INM-7: Brain and Behavior), Research Centre Jülich, Jülich, Germany.

**Keywords:** Asymmetry, Lateralization, Hemispheric difference, Functional gradients, Macaques, Evolution, Large-scale organization, Cerebral cortex, Heritability

## Abstract

The human cerebral cortex is symmetrically organized along large-scale axes but also presents inter-hemispheric differences in structure and function. The quantified contralateral homologous difference, i.e., asymmetry, is a key feature of the human brain left-right axis supporting functional processes, such as language. Here, we assessed whether the asymmetry of cortical functional organization is heritable and phylogenetically conserved between humans and macaques. Our findings indicate asymmetric organization along an axis describing a hierarchical functional trajectory from perceptual/action to abstract cognition. Whereas language network showed leftward asymmetric organization, frontoparietal network showed rightward asymmetric organization. These asymmetries were heritable and comparable between humans and macaques, suggesting (phylo)genetic conservation. However, both language and frontoparietal networks showed a qualitatively larger asymmetry in humans relative to macaques and variable heritability in humans. This may reflect an evolutionary adaptation allowing for experience-dependent specialization, linked to higher-order cognitive functions uniquely developed in humans.

## Introduction

The human cerebral cortex consists of two hemispheres that are not exactly alike and show marked differences in structure and function along a left-to-right axis ^1–9^. It has been suggested that the brain favors asymmetry to avoid duplication of neural circuitry having equivalent functions ^6, 10^. For example, bilateral cortical regions showing asymmetry in task-evoked activity have reduced (long-range) connections with the opposite homologous regions, favoring more local connectivity ^6^.

Asymmetry, i.e., quantitative hemispheric differences between contralateral homologous regions, supports partly differentiable functional processes ^6, 11, 12^. Previous work has suggested that functions related to leftward dominance include language processing ^13–15^, letter search ^16^, and analogical reasoning ^17^. On the other hand, rightward dominance of functional activation has been related to holistic word processing ^18^, visuospatial abilities ^19^, emotional processing ^20^, as well as with psychiatric disorders such as autism spectrum disorder ^21^. In addition to task-related asymmetries, resting state functional connectivity (FC) studies have also reported hemispheric differences. For example, language areas of the middle and superior temporal cortex showed increased connectivity with regions in the left hemisphere relative to their right hemispheric counterparts ^4^, and the right amygdala showed higher connectivity with the entire cortex than the left amygdala ^22^. Moreover, previous work has indicated that there are inter- and intra-hemispheric differences in functional connectivity between healthy adults and patients with schizophrenia ^23^, and between neurotypical individuals and those diagnosed with autism spectrum disorder ^24^. It is possible that such functional processing asymmetries may be driven by subtle differences in functional organization between the hemispheres.

One appealing approach to studying functional organization is by evaluating the low-dimensional axes, or gradients, present within the connectome. These approaches embed brain regions on a continuous data-driven space based on their functional connectome ^25–27^. Gradients capture how connectivity profiles from distinct cortical regions are integrated (i.e. similar functional connectivity profiles) and segregated (i.e. dissimilar functional connectivity profiles) across the cortex ^25, 28–30^. Regions that have similar connectivity profiles are at similar positions along these gradients, whereas regions with dissimilar connectivity profiles are placed further apart. The principal functional gradient, partly reflected in the intrinsic geometry of the cortex, shows that regions of the transmodal systems occupy locations equidistant from unimodal systems ^25, 31, 32^. This pattern describes a functional hierarchy from perceptual/action to more abstract cognitive processes. Moreover, cortical gradients describe variations in genetic patterning ^33, 34^, functional processes ^25, 32, 35^, and are observed across species ^33, 36, 37^. Notably, recent research suggests that the principal gradient is asymmetric ^9, 38^ and that the degree of asymmetry relates to individual differences in semantic performance and visual reasoning ^38^.

As inter-hemispheric asymmetry has been observed consistently in human brain structure and function, there may be important (phylo)genetic factors supporting lateralized human cognition ^2, 39–43^. Previous work has reported that brain structure asymmetry is heritable ^1, 3^, especially in the language areas, and differentiates between humans and non-human primates ^44–47^. At the same time, it has been shown that both humans and apes show asymmetry of brain shape ^47^, indicating that asymmetry is not a uniquely human brain feature. However, asymmetry was observed to be more local and variable in humans, potentially suggesting that individual variation in asymmetry in humans varies as a function of localized networks rather than global features.

Here, we investigated the genetic basis of asymmetry of functional organization. We first examined whether inter-individual differences in asymmetry of functional organization are under genetic control, i.e., heritable. Second, we investigated whether asymmetry of functional organization is phylogenetically conserved in macaques. To probe individual variation in asymmetry of functional organization we utilized a data-driven nonlinear dimension reduction technique, as this approach can provide reliable and robust indices of individual variation of cortical organization ^31^. We first obtained connectomic gradients for each hemisphere separately (left and right intra-hemispheric) as well as for both hemispheres together (left and right inter- hemispheric). Subsequently, to evaluate the heritability of possible differences between left and right intra- and inter-hemispheric FC gradients, we used the twin pedigree set-up of the Human Connectome Project S1200 release young adults dataset ^48^. To assess whether asymmetry is conserved in other primates, we compared the asymmetry of functional gradients of humans with those observed in macaque monkeys using the prime-DE dataset ^36, 49^. Finally, we conducted a confirmatory meta-analysis to explore the relationship between the patterns of gradient asymmetry and task-based functional MRI activations. Multiple analyses verified the robustness and replicability of our results.

## Results

### Hemispheric functional connectivity gradients (Figure 1)

To obtain intra-hemispheric gradients, we first computed the functional connectivity (FC) in 180 homologous parcels per hemisphere using a multimodal parcellation (MMP, ^50^ for each subject (n = 1014). For the network level analyses, we employed the Cole-Anticevic atlas ^51^ based on the MMP (**Fig. 1a)**. For each individual, FC was summarized in two different patterns (**Fig. 1b**): FC within the left hemisphere (LL mode, intra-hemispheric pattern), within the right hemisphere (RR mode, intra-hemispheric pattern), from left to right hemisphere (LR mode, inter-hemispheric pattern), and from right to left hemisphere (RL mode, inter-hemispheric pattern). We selected the LL mode as the reference template for the gradients approach, and therefore assessed the mean FC that was determined by averaging LL FC across subjects (lower panel in **Fig. 1c**). Here, the reference matches the order and direction of the gradient but does not rescale the gradients. The template gradients were computed by implementing diffusion map embedding, a non-linear dimension reduction technique ^27^, on the mean LL FC using BrainSpace ^26^. The first three gradients (G1, G2, G3) explained approximately 23.3%, 18.1%, and 15.0% of the total variance in the LL functional connectome (**Fig. 1d)**.

**Fig. 1.**
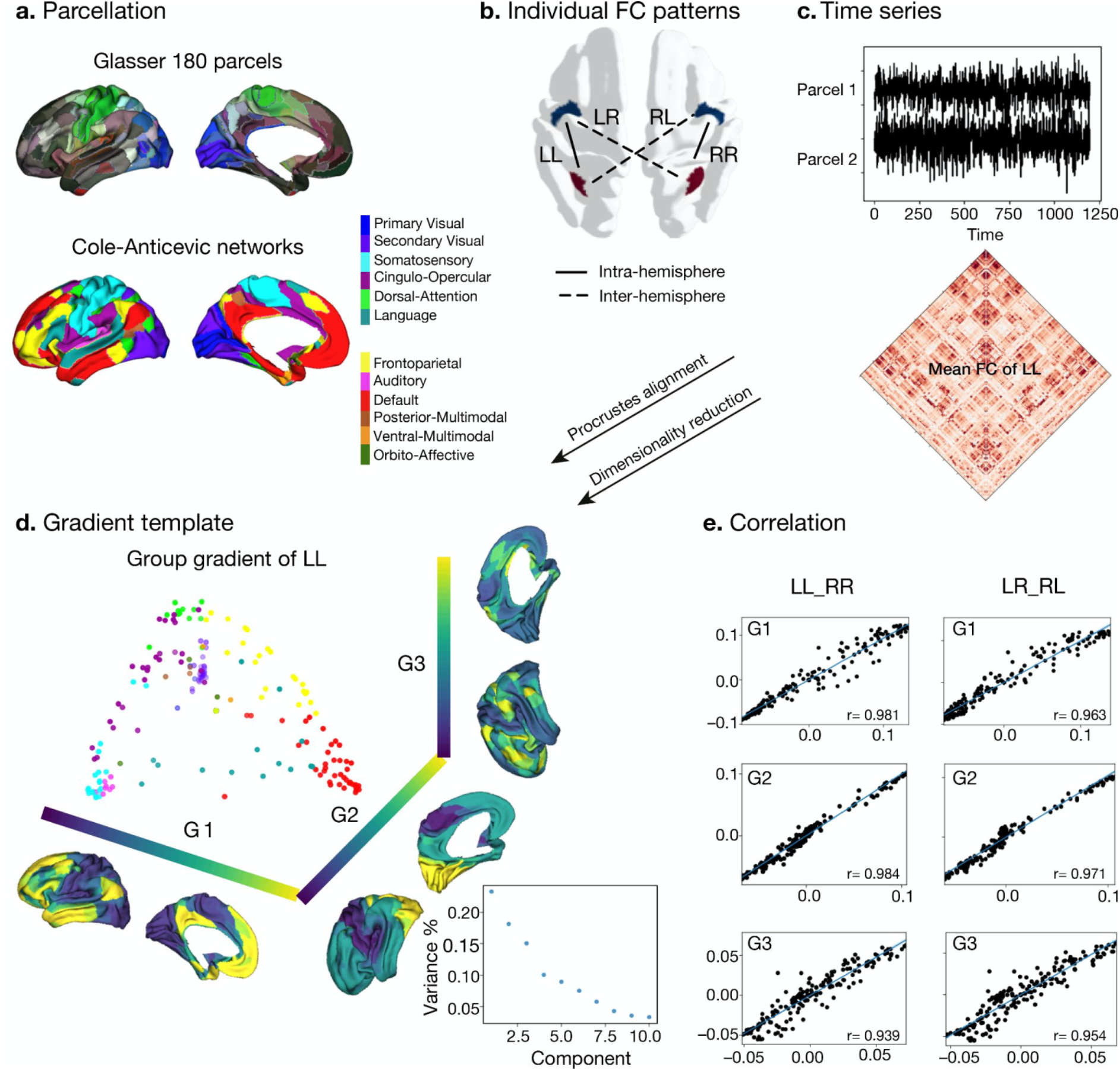
Processing of functional gradients in humans. **a)** Parcellation using Glasser 180 atlas ^50^ in each hemisphere and Cole-Anticevic networks ^51^ for humans. **b)** Individual FC in each hemispheric pattern, i.e., left-left (LL, intra-hemisphere), right-right (RR, intra-hemisphere), left-right (LR, inter-hemisphere), and right-left (RL, inter-hemisphere). **c)** Time series of two parcels and the mean functional connectivity (FC) matrix between left and left hemisphere (LL). **d)** Gradient template using the group-level gradient of LL. Dots represent parcels and were colored according to Cole-Anticevic networks. **e)** Correlation between left and right mean gradients across subjects of intra- and inter-hemispheric patterns.

Next, individual gradients were computed for each subject and the four different FC modes and aligned to the template gradients with Procrustes rotations. As noted, Procrustes matching was applied without a scaling factor so that the reference template only matters for matching the order and direction of the gradients. **Fig.1e** shows the correlation between LL and RR, LR, and RL modes. In each case the gradients were highly similar. Similar to previous work (Margulies, 2016) we observed that the principal gradient (G1) traversed between unimodal regions and transmodal regions (e.g. default-mode network: DMN) whereas a visual to somatosensory gradient was found for G2. The tertiary gradient (G3) dissociated control from DMN and sensory-motor networks (**Fig. 1d and 1e, and Supplementary Fig. S1**). We employed generative modeling of brain maps with spatial autocorrelation ^52^ for correcting Spearman correlation *P* values, i.e., *P* _gen_. For the intra-hemispheric pattern, the mean gradients of LL were strongly correlated with those of RR (Spearman *r*_G1_ = 0.981, *P* _gen_ < 0.001, *r*_G2_ = 0.984, *P* _gen_ < 0.001, *r*_G3_ = 0.939, *P* _gen_ < 0.001). For the inter-hemispheric pattern, the mean gradients of LR were also strongly correlated with those of RL (Spearman *r*_G1_ = 0.963, *P* _gen_ < 0.001, *r*_G2_ = 0.971, *P* _gen_ < 0.001, *r*_G3_ = 0.954, *P* _gen_ < 0.001).

### Asymmetry of functional gradients in humans (Figure 2)

Next, we computed the asymmetry index (AI) by subtracting the left hemispheric gradient scores of each parcel from the corresponding right hemispheric scores for our intra- and inter- hemispheric connectivity patterns (**Fig. 2a**). A red AI indicates rightward dominance in gradient scores, whereas blue indicates leftward dominance. The significance of AI scores for the intra- and inter-hemispheric patterns were reported after false discovery rate adjustment (*P* _FDR_ < 0.05) (**Fig. 2b**). Frontal and temporal lobes showed the greatest intra-hemispheric asymmetry in G1 (**Supplementary Table S1**). In particular, regions in ventral- and dorsolateral PFC (11l, p9-46v, 9-46d; ^50^) were the three most rightward asymmetric areas and regions in temporal polar cortex, dorso/posterior superior temporal sulcus, and dorsolateral PFC (TGv, STSdp, and 55b) were the three most leftward asymmetric areas in the intra-hemispheric pattern. Network-level analyses (**Fig. 2c)** indicated that the language (*t* = 41.3, *df* = 1013, *P* _FDR_ < 0.001) and default mode (*t* = 17.3, *df* = 1013, *P* _FDR_< 0.001) networks had a high leftward AI, while the frontoparietal network (*t* = -26.0, *df* = 1013, *P* _FDR_ < 0.001) had a high rightward AI. We observed no significant difference of AI in primary and secondary visual networks. Overall, asymmetry was widely present along the first three connectivity gradients, including G2 and G3 (**Supplementary Fig. S2**).

**Fig. 2.**
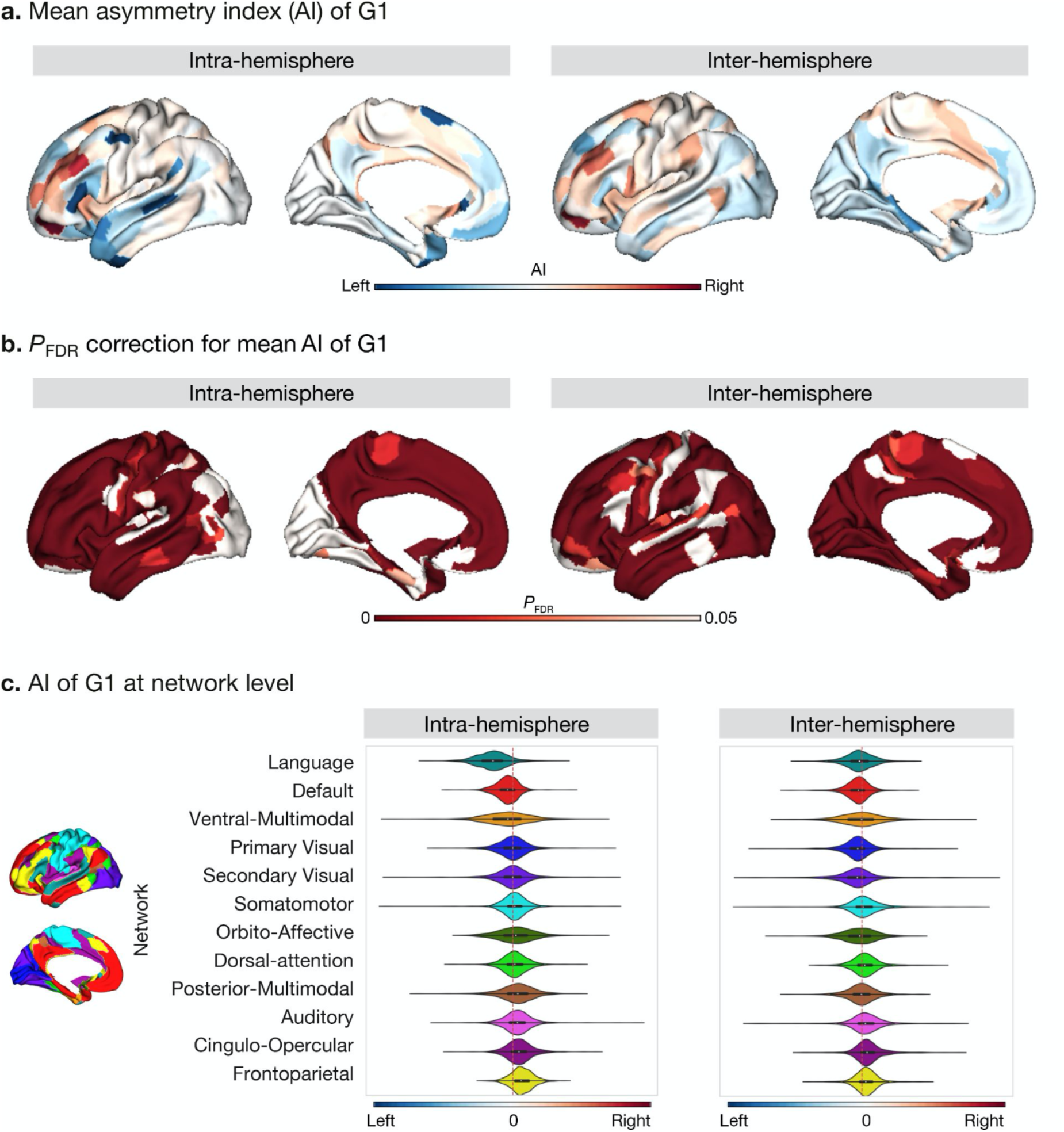
Asymmetry of functional gradients in humans and its heritability. **a)** Mean asymmetry index (AI) of intra-hemispheric and inter-hemispheric patterns in humans. Red indicates rightward preference and blue indicates leftward preference. **b)** FDR correction for the *P* value of AI shown in A. **c)** Asymmetry of G1 at the network level using Cole-Anticevic networks ^51^.

For the inter-hemispheric pattern, a large portion of the cerebral cortex showed significant AI scores. The top six asymmetric areas included regions in inferior frontal cortex and parahippocampal regions (11l, 46, 9-46v, p9-46v, PreS, and PHA2) (**Supplementary Table S1**). At the network level (**Fig. 2c)**, networks with leftward dominance were the visual (*t* _primary visual_ = 9.3, *df* = 1013, *P* _FDR_ < 0.0019; *t* _secondary visual_ = 7.5, *df* = 1013, *P* _FDR_ < 0.001), language (*t* = 5.7, *df* = 1013, *P* _FDR_ < 0.001), default mode (*t* = 11.9, *df* = 1013, *P* _FDR_ < 0.001), and orbito-affective (*t* = 4.6, *df* = 1013, *P* _FDR_ < 0.001) networks. Networks with rightward dominance were the somatomotor (*t* = -3.5, *df* = 1013, *P* _FDR_ < 0.00), cingulo-opercular (*t* = -14.6, *df* = 1013, *P* _FDR_ < 0.001), dorsal attention (*t* = -8.0, *df* = 1013, *P* _FDR_ < 0.001), frontoparietal (*t* = -12.0, *df* = 1013, *P* _FDR_ < 0.001), and auditory (*P* _FDR_ < 0.001) networks. Posterior and ventral multimodal networks were not significantly asymmetric.

The mean AI scores along G1 across individuals for the intra- and inter-hemispheric patterns showed high similarity (Spearman *r* = 0.710, *P* _gen_ < 0.001). This may indicate that the asymmetric functional organization is a feature that is captured both by inter- and intra-hemispheric connectivity patterns.

### Heritability of asymmetry of functional gradients in humans (Figure 3)

We next computed the heritability of the AI scores of the functional gradient for the intra- and inter-hemispheric patterns using Solar-Eclipse 8.5.1 beta (http://solar-eclipse-genetics.org/). We found that left-right differences observed in large-scale functional organization axes were heritable (**Fig. 3a**). Specifically, for the intra-hemispheric pattern, we found sensory-motor regions, middle temporal regions, dorso-lateral, and medial prefrontal regions to be heritable (*P* _FDR< 0.05_). In the case of the inter-hemispheric pattern, all cortical regions with the exception of visual areas and superior temporal and insular regions were heritable (*P* _FDR< 0.05_). Notably, language-associated areas such as the PSL (Peri-Sylvian language area) and 55b had the highest heritability in both the hemispheric patterns (PSL: intra: *h*^2^ = 0.39, *P*_FDR_ < 0.001 and inter: *h*^2^ = 0.30, *P* _FDR_ < 0.001, **Supplementary Table S1**). However, area 44 (Broca’s area) showed low heritability (intra: *h*^2^= 0.14, *P*_FDR_ = 0.026 and inter: *h*^2^= 0.16, *P*_FDR_ = 0.018). The G2 and G3 results are shown in **Supplementary Fig. S2**.

**Fig 3.**
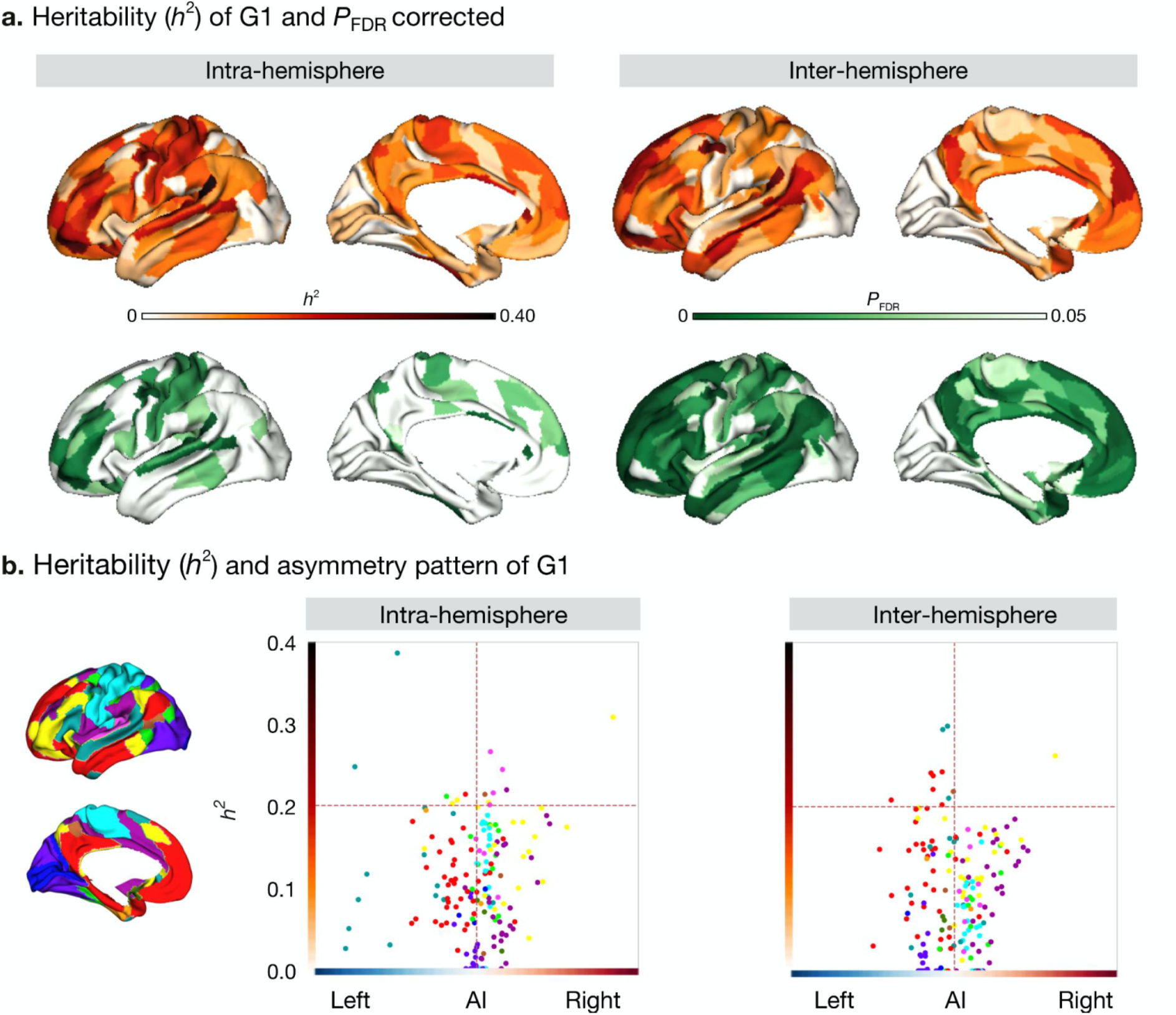
Heritability of asymmetry of functional G1. **a**) Heritability (orange colorbar) and *P* values after FDR correction (green colorbar). **b**) Scatter plot of heritability and AI scores. Dots represent parcels and are colored according to CA networks ^51^.

To assess whether regions showing higher asymmetry had an increased heritability of G1, we correlated our cortical maps of asymmetry with those reporting heritability (**Fig. 3b**). For the correlation between the absolute asymmetry index and heritability, gradients of the intra-hemispheric FC patterns were significant (*r* = 0.321, *P* _gen_ = 0.009) but gradients of the inter- hemispheric FC were not (*r* = 0.082, *P* _gen_ = 0.641).

### Asymmetry of functional gradients in macaques (Figure 4)

To probe the phylogenetic conservation of asymmetry of functional organization in primates, we performed the same diffusion map embedding analysis on macaque resting-state FC data (n = 19, PRIMATE-DE sample, ^36, 49^. We used the Markov parcellation ^53^ in macaques, resulting in 91 parcels per hemisphere (**Fig. 4a**) and then computed FC in the four patterns: LL and RR (intra-hemispheric patterns), and LR and RL (inter-hemispheric patterns). Following the same connectome gradients analysis pipeline as deployed on the human FC data, we obtained the template gradients on the LL intra-hemispheric FC pattern (**Fig. 4b**). The first three template gradients explained 18.9%, 15.1%, and 12.8% of total variance respectively. G1 described an axis traversing dorsolateral prefrontal and parietal regions (anterior-posterior).

**Fig. 4.**
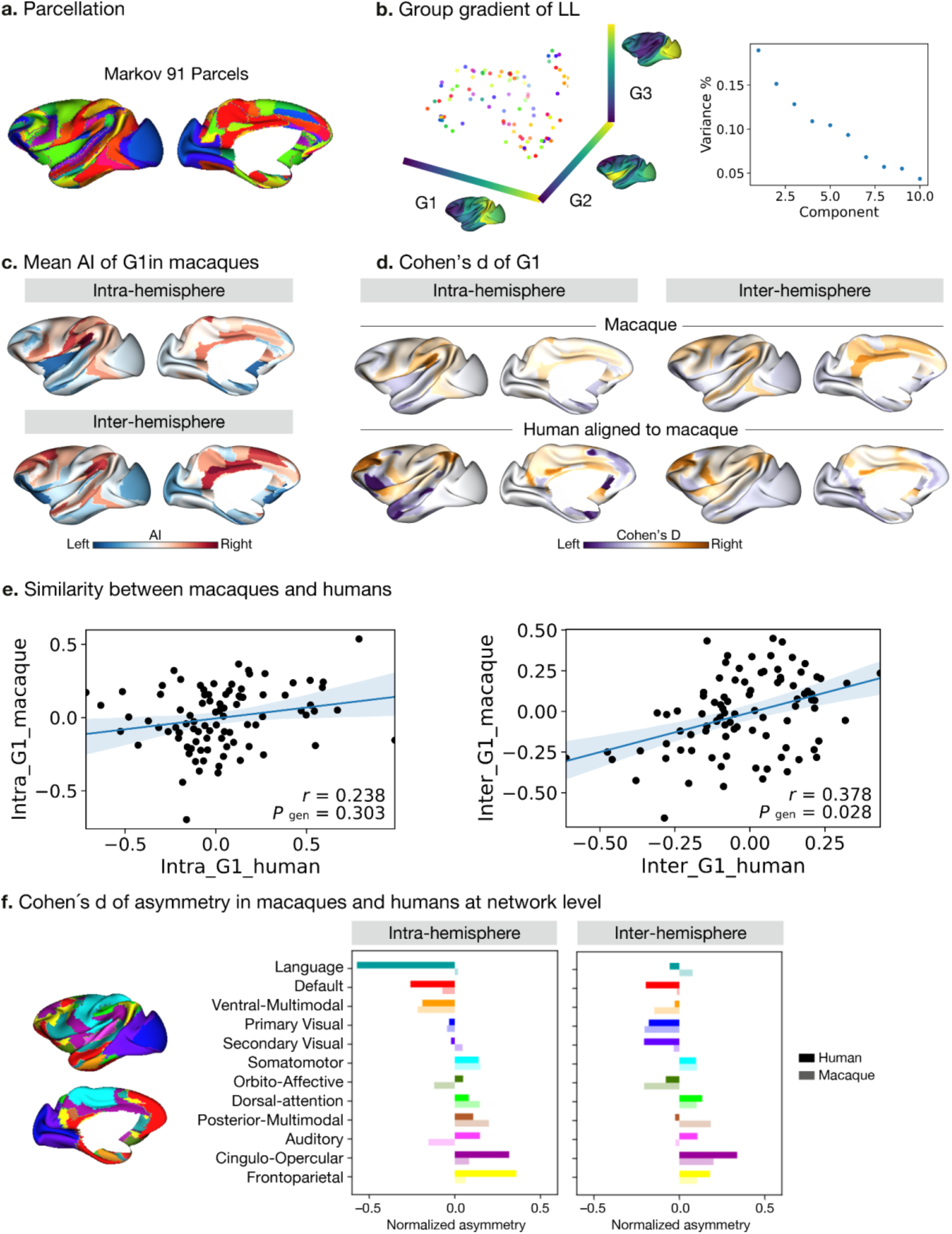
Asymmetry of functional gradients in macaques. **a)** Parcellation used Markov 91 atlas in macaques ^53^. **b)** Template gradients of group level connectivity of LL. **c)** Mean asymmetry index of G1 in macaques. **d)** Normalized (Cohen’s d) asymmetry of G1 in macaques and humans aligned to macaque’s surface. **e**) Similarity of normalized asymmetry of G1 between humans and macaques. **f)** Comparison between macaques and humans in Cohen’s d of asymmetry of G1. The projection of human Cole-Anticevic networks ^51^ to the macaque surface can be seen in Methods for details. Bold colors indicate human results and muted colors indicate macaque results.

Evaluating the intra-hemispheric pattern of functional organization in macaques along G1, we observed that parietal cortices had a rightward dominance while occipital cortices were leftward. Temporal cortex asymmetry was low (**Fig. 4c**). The inter-hemispheric pattern showed similar asymmetry to the intra-hemispheric pattern along G1. However, the AI scores of the principal, but also secondary and tertiary gradients, were not statistically significant after FDR correction, both for intra- and inter-hemispheric patterns. The effect sizes across cortex observed in macaques along G1 were [intra: mean Cohen’s d = -0.18 (rightward) and 0.16 (leftward); inter: mean Cohen’s d = -0.20 (rightward) and 0.18 (leftward)].

To compare human and macaque connectomic gradients, we aligned human gradients to the same macaque surface space (**Fig. 4d**) using a joint embedding technique ^36^. We summarized Cohens’ d of AI of macaque-aligned human gradients within the Markov parcels for the intra- and inter-hemispheric patterns and compared the similarity of Cohens’ d of AI between the two species using Spearman correlations (**Fig. 4e**). We found that the macaque and macaque- aligned human AI maps of G1 were correlated positively for inter-hemispheric patterns (*r* _inter- hemisphere_ = 0.378, *P* _gen_ = 0.028). For intra-hemispheric patterns, we only observed a positive association at uncorrected level (*r* _intra-hemisphere_ = 0.238, *P* _gen_ = 0.303, *P* _uncorrected_ = 0.023). Exploring potential correlations between the first three gradients of macaques and humans (**Supplementary Fig. S3**), we found significant positive correlation between inter- macaque G1 and human G2 (*r* = 0.378, *P* _gen_ = 0.034), and negative correlation between intra- macaque G2 and human G1 (*r* = -0.607, *P* _gen_ = 0.001), and inter macaque G3 and human G1 (*r* = -0.480, *P* _gen_ < 0.001).

We then projected the human functional networks ^51^ on the macaque surface ^36^, to qualitatively compare differences in human functional networks between humans and macaques (**Fig. 4f)**. In the case of the intra-hemispheric asymmetry of the principal FC gradient, we observed that humans showed high leftward asymmetry in the language and default mode networks but macaques did not. Moreover, humans showed high rightward asymmetry in the frontoparietal and cingulo-opercular networks but macaques did not. Humans and macaques showed an opposite direction of asymmetry in auditory, orbito-affective, and secondary visual networks. For the inter-hemispheric FC pattern, macaques and humans showed only subtle differences.

### Functional decoding along the normalized asymmetry of G1 (Figure 5)

Finally, we investigated the relationship between patterns of asymmetry of functional organization in humans and task-based meta-analytic functional activations. To do so, we projected meta-analytical fMRI activation maps ^54^ along the normalized (Cohen’s d) asymmetry of G1 (**Fig. 5**). Our choice for the 24 cognitive domain terms were consistent with prior literature ^25^. Here, we calculated the weighted score by activation z-score (parcels where activation z-score was greater than 0) multiplied by the normalized asymmetry, suggesting leftward to rightward preference, seen from top to bottom of the y-axis of **Fig. 5**. Language, autobiographical memory, and social cognition domains were associated with leftward hemispheric preference, whereas cognitive control, working memory, and inhibition were associated with rightward hemispheric preference. For the asymmetry of the inter-hemispheric FC gradient, we observed a similar pattern of association (**Supplementary Fig. S5**). This indicates that patterns of asymmetry in functional organization also align with task-based activations consistently reported in the literature.

**Fig. 5.**
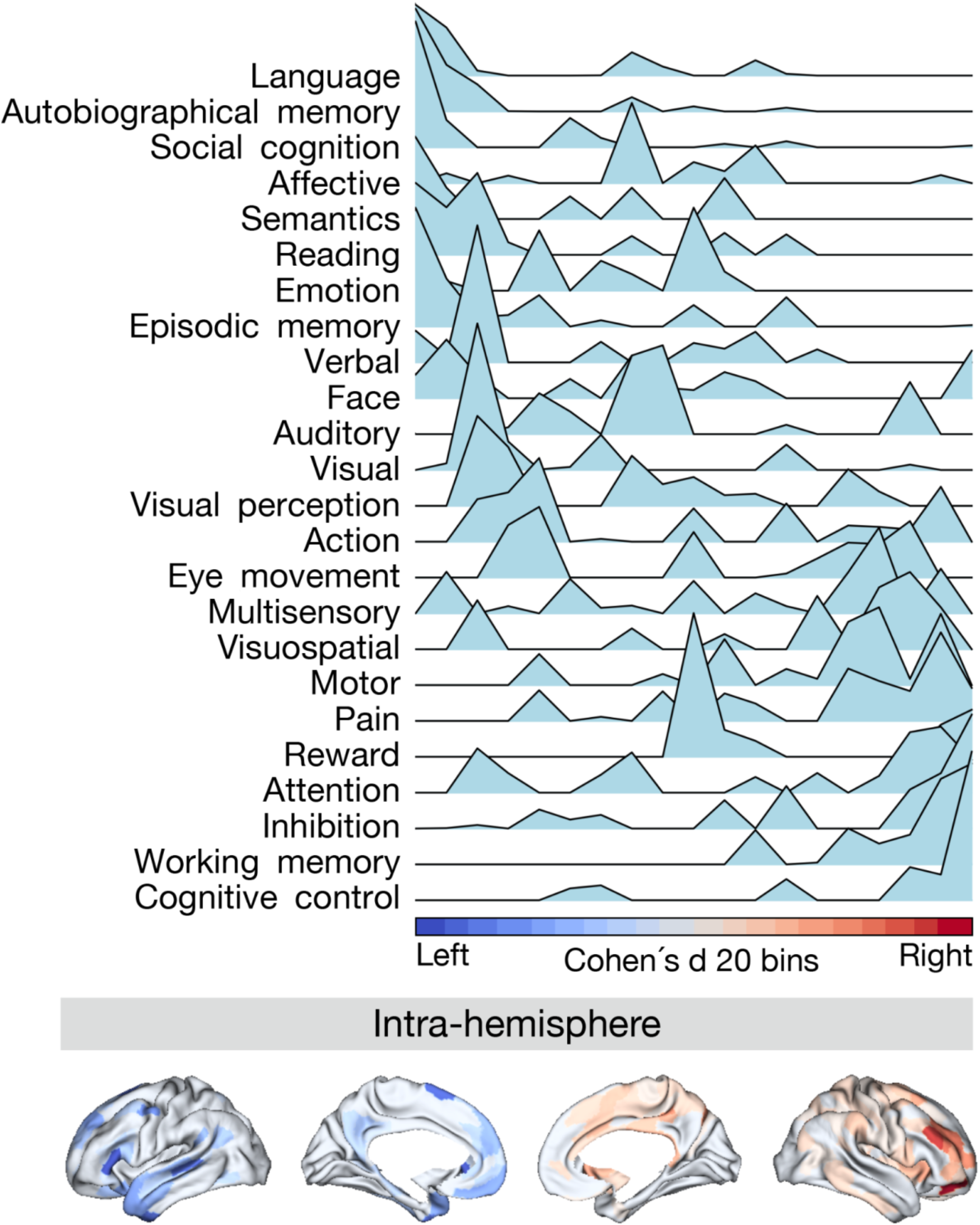
Projection of meta-analytical task-based function along normalized asymmetry of G1 (intra-hemisphere). The 20 bins were generated by normalized (Cohen’s d) asymmetry of G1 in humans. Cool color indicates regions showing leftward dominance and warm color indicates regions showing rightward dominance. The order of the terms of the y-axis was generated by the weighted score of activation (z-score > 0) * normalized asymmetry.

### Robustness analyses

To determine the robustness of our results, we performed three distinct manipulations. First, we examined AI using the normalized formula ^5^, i.e., (left-right)/(left+right). Secondly, we tried a different parcellation scheme, i.e. the Desikan-Killiany (DK) atlas (34 cortical parcels per hemisphere, ^55^. This scheme has commonly been used in previous literature on asymmetry ^1, 5, 7^. Finally, we tested another large human data sample (UK Biobank, UKB dataset n = 34,604). Overall, we observed that for human G1, the results were robust using the normalized AI formula (**Fig. S6**). The DK atlas results were similarly able to detect asymmetric effects in large-scale parcels, but given the reduced spatial resolution not the subtle differences (**Fig. S7**). For the G1 of the intra-hemispheric FC pattern, we observed a correlation between UKB and HCP samples (Spearman *r* = 0.269, *P* _gen_ = 0.022). However, associations between HCP and UKB were not significant for the inter-hemispheric asymmetry pattern of G1 (Spearman *r* = 0.043, *P* _gen_ = 0.852) (**Fig. S8**). Both HCP and UKB showed most leftward asymmetry in language and default mode networks but rightward frontoparietal network in HCP was not observed in UKB. Detailed results are reported in the **Supplementary Materials**.

## Discussion

In this study, we investigated the extent to which human cortical functional organization is asymmetric using a gradient-based approach. We assessed whether genetic factors shape such asymmetry and evaluated whether patterns of asymmetry are phylogenetically conserved between humans and non-human primates (macaques). We found that the principal gradient revealed hemispheric differences in most cortical regions, excluding the visual cortex. The language network and default-mode network showed the most leftward asymmetry while the frontoparietal network showed the most rightward asymmetry. The observed asymmetry of functional organization along the principal gradient was heritable. At the same time, regions with high asymmetry showed variable heritability. This may suggest that asymmetry in functional organization reflects both heritable and experience-dependent factors. Although the difference in left and right hemispheric functional organization was not significant along the principal functional gradient in a sample of macaques, in particular the inter-hemispheric asymmetric pattern was comparable to the asymmetry pattern observed in humans indicating phylogenetic conservation. Notably, both the language and frontoparietal networks showed a higher leftward asymmetry in humans relative to macaques, indicating cross-species differences in asymmetry of specific transmodal functional networks. Decoding task-based functional activations along the asymmetry axis of the principal gradient, we observed that regions with a leftward preference were associated with language, autobiographical memory, and social cognition domains, whereas those with a rightward preference included cognitive control, working memory, and inhibition. In sum, our study shows the asymmetry of functional organization is, in part, heritable in humans and phylogenetically conserved in humans and macaques. At the same time we observed that asymmetry of regions linked to higher-order cognitive functions such as language and cognitive control showed marked differences between humans and macaques and variable heritability in humans, possibly reflecting an evolutionary adaptation allowing for experience-dependent specialization.

By studying asymmetry in functional organization using a gradient approach, we have extended previous studies reporting asymmetric functional connectivity. Indeed, although the functional organization of the cerebral cortex has a largely symmetric pattern, it also shows subtle differences between hemispheres ^9, 38, 56, 57^. For the intra-hemispheric asymmetry gradients, we found that regions belonging to the language network showed the strongest leftward preference along the principal gradient axis. This indicates that their functional connectivity profiles were more similar to the default mode, relative to their right-hemispheric counterparts. Conversely, ventral multimodal networks were closer to the transmodal apex of the principal gradient in the right hemisphere, relative to their homologues in the left hemisphere. As such, our observations suggest that key transmodal regions, part of the language and control networks, show organizational preference to either the left or right hemisphere. Anterior lateral default mode subnetworks have been shown to uniquely exhibit positive connectivity to the language network ^58^, possibly leading to increased gradient loadings of the language network in the left hemisphere, placing them closer to the default regions along the principal gradient in the left hemisphere relative to the right. Conversely, the transmodal frontoparietal network was located at the apex of rightward preference, possibly suggesting a right-ward lateralization of cortical regions associated with attention and control and ‘default’ internal cognition ^59, 60^. The observed dissociation between language and control networks is also in line with previous work suggesting an inverse pattern of language and attention between hemispheres ^6, 61^. Such patterns may be linked to inhibition of corpus callosum ^62^, promoting hemispheric specialization. In the current study, we found highly similar patterns when comparing differences in inter- and intra-hemispheric functional connectivity, possibly suggesting both are closely coordinated in young adulthood. It has been suggested that such inter-hemispheric connections set the stage for intra-hemispheric patterns related to association fibers ^63^. Future research may relate functional asymmetry directly to asymmetry in underlying structure to uncover how different white-matter tracts contribute to asymmetry of functional organization.

We furthermore investigated whether such individual variations in asymmetry of functional organization could be attributed to genetic factors. To do so, we performed heritability analysis enabled by the twin design of the Human Connectome Project ^48^. Previous work indicated that brain structure including cortical thickness, surface area, and white matter connection ^1, 3, 7^, as well as functional connectome organization ^64, 65^ are heritable. Our twin-based heritability analyses revealed heritable asymmetry of the principal functional gradient in the entire cortex, excluding visual cortex. At the same time, studying the association between heritability and asymmetry patterns we observed mixed results. Though we observed that the language-related area PSL showed the highest heritability, the highly asymmetric area 44 (Broca’s area) showed the lowest heritability. This may reflect a differential (dorsal and ventral) pathway of language development in the frontal and temporal lobe, where the dorsal pathway to the inferior frontal gyrus matures at later stages in development ^66^. For example, previous work found that temporal language areas showed high heritability of cortical thickness asymmetry ^1^ and white matter connection asymmetry ^3^ but frontal language areas did not. Such posterior-anterior differences may be due to developmental factors or axes of stability versus plasticity in the cortex ^67^. A case study of an individual born without a left temporal lobe found that frontal language areas in the left hemisphere did not emerge in the absence of temporal language areas in the left hemisphere, and that language functions instead relied on the right hemispheric functional network ^68^. It is thus possible that Broca’s area may mature after more posterior language regions in hierarchical fashion, which may be related to decreasing heritability in frontal language areas (i.e., more influenced by developmental and/or environmental factors). Recent work suggests that asymmetric patterning of brain structure and function are largely determined prenatally and unaffected by preterm birth ^69^. In neonates asymmetric patterns were largely observed in primary and unimodal areas, whereas association regions were largely symmetric. Thus, asymmetry in association regions may be more experience-dependent, reflected in our study by variable heritability. One focus of future work could thus be to evaluate the development of asymmetry in functional organization. Moreover, by means of GWAS approaches, it may also be possible to get more insight in specific genes and associated processes involved in functional asymmetry.

Evaluating the correspondence of asymmetry of functional organization between humans and macaques, by aligning the human gradients to the macaque gradient space ^36^, we observed some similarity between asymmetry of the gradients in both species. This indicates functional asymmetry may be, in part, conserved across primates. At the same time, we found that language, default mode, and frontoparietal networks showed qualitatively more asymmetry in humans (human > macaque). These findings may support the notion that though asymmetry is a phenomenon existent across different primates, regions involved in higher-order cognitive functions in humans are particularly asymmetric. Previous work studying asymmetry in white matter structure in primates found that humans showed more leftward arcuate fasciculus volume and surface relative to macaques ^44^. The arcuate fasciculus is a white matter tract implicated in language functions by connecting Broca’s and Wernicke’s areas ^70^. Moreover, by comparing humans, macaques, and chimpanzees, ^45^ report evolutionary modifications to this tract in humans relative to other primates, possibly derived from auditory pathways ^71^. At the same time, other structural studies have also observed leftward asymmetry of language areas in chimpanzees, indicating that asymmetry of language-regions per se may not be a human- specific feature ^46, 72^. Fittingly, there are no significant differences of thickness and area asymmetry between humans and chimpanzees in superior temporal lobe ^72^. Studying the endocranial shape of humans and non-human primates, temporal and occipital cortices showed local differences in asymmetry across species, and much more variability in humans relative to non-human primates ^47^. This suggests that whereas brain asymmetry is a phenomenon observed throughout mammals ^73^, specific nuances may relate to species-specific behavioral and cognitive differences. Future research could assess asymmetry of brain organization in other primates, and relate asymmetry of brain structure and function to behavioral differences across primates.

The functional relevance of asymmetry along the sensory-transmodal axis was evaluated in the human brain by projecting meta-analytical task-based coactivations along asymmetric effects of the functional principal gradient. In line with our expectations based on the distribution of asymmetry within functional networks, we found that task-based activations associated with language processing leaned leftward while task-based activations associated with executive functions leaned rightward, specifically in the intra-hemispheric pattern. This suggests that lateralized functions supported by the brain’s asymmetry have functional relevance (especially higher-order cognitive functions such as language and executive control). Indeed, related work has shown a direct link between asymmetry and semantic and visual recognition skills ^38^, suggesting that asymmetry of individuals relates to variation in behavioral performance in these domains ^38, 74^. Our observation of asymmetry of language versus executive functions may also be in line with notions of differential axes of asymmetry, dissociating symbolic/language, emotion, perception/action, and decision functional axes ^6^. The asymmetry of principal functional gradient in humans and macaques showed a divergence along these axes, possibly indicating cross-species variability within the lateralization archetypes in primates. Notably, left hemispheric language lateralization is enabled throughout language development while right hemispheric language activation declines systematically with age ^75^. Therefore, future research may focus on studying how the lateralization of human behavior is shaped by development and aging and how this may impact function and behavior.

Though we showed asymmetry in functional organization, there are various technical and methodological aspects to be considered. In the current work we used the MMP ^50^ for surface-based human fMRI data. A previous study used the atlas of intrinsic connectivity of homotopic areas ^76^, AICHA, www.gin.cnrs.fr/en/tools/aicha) for voxel-based fMRI data ^9^. In line with the results of that study, we found similar intra-hemispheric differences in functional gradients. Extending that work we additionally used the DK atlas ^55^, which is often used in structural asymmetry studies ^1, 77^. We again found asymmetric patterns, with a rightward dorsal frontal lobe and leftward posterior superior temporal lobe. The other temporal regions, having leftward or rightward asymmetry using MMP ^50^, showed no or less asymmetry using the DK atlas. Possibly, such subtle differences are not captured by the DK atlas, with only 34 cortical parcels per hemisphere. Evaluating the consistency of functional asymmetry across different datasets, we found that HCP (n = 1014) and UKB (n = 34,604) showed consistent leftward asymmetric functional organization in the language and default mode networks but no consistent rightward asymmetry of the frontoparietal network. Such differences may be due to technical differences between the datasets ^78^. However, it may also reflect sample specific differences in asymmetry. Indeed, whereas the HCP sample consists of young-adults with an age-range of 22 - 37 years, the UKB has a comparatively older and wider age range (from 40 to more than 70 years). Thus, it is possible the observed differences in the frontoparietal network are directly related to age-related asymmetry effects ^75^.

To conclude, we investigated the genetic and phylogenetic basis of asymmetry of large-scale functional organization. We observed that the principal (unimodal-transmodal) gradient ^25^ is asymmetric, with regions involved in language showing leftward organization and regions associated with executive function showing rightward organization. This asymmetry was, in part, heritable and phylogenetically conserved. However, functional asymmetry was more pronounced in language networks in humans relative to macaques. The current framework may be expanded by future research investigating the development and phylogeny of functional asymmetry as well as its neuroanatomical basis in healthy and clinical samples. This may provide important insights in individual-level brain asymmetry and its relation to human cognition.

## Methods

### Participants

#### Humans

For the analyses in humans, we used the Human Connectome Project (HCP) S1200 data release ^48^. That release contains four sessions of resting state (rs) fMRI scans for 1206 healthy young adults and their pedigree information (298 monozygotic and 188 dizygotic twins as well as 720 singletons). We included individuals with a complete set of four fMRI scans that passed the HCP quality assessment ^48, 79^. Finally, our sample consisted of 1014 subjects (470 males) with a mean age of 28.7 years (range: 22 - 37).

The rs-fMRI data was collected at two sessions and in two phase encoding directions at each session (left-right [LR1], [LR2] and right-left [RL1], [RL2]). All rs-fMRI data underwent HCP’s minimal preprocessing ^79^ and were coregistered using a multimodal surface matching algorithm (MSMAll) ^80^ to the HCP template 32k_LR surface space. The template consists of 32,492 total vertices per hemisphere (59,412 excluding the medial wall).

For the replication, we employed the UKB dataset (application ID: 41655) including 34,604 subjects’ imaging data. Details on data processing and acquisition can be found in the UKB Brain imaging documentation (https://biobank.ctsu.ox.ac.uk/crystal/crystal/docs/brain_mri.pdf). Briefly, resting-state imaging data was motion corrected, intensity normalized, high-pass temporally filtered, and further denoised using the ICA-FIX pipeline, all implemented in FSL. MPM parcellation was warped to subject-space based on the high-resolution T1-weighted anatomical image. Individual warping parameters were applied to map the MPM parcellation to the functional space following T1-rsfMRI alignment. The age range of the UKB sample was from 40 to more than 70 years.

#### Macaques

We selected rhesus macaque monkeys’ rs-fMRI data from the non-human primate (NHP) consortium PRIME-DE (http://fcon_1000.projects.nitrc.org/indi/indiPRIME.html) from Oxford. The full dataset consisted of 20 rhesus macaque monkeys (macaca mulatta) scanned on a 3T with a 4-channel coil ^81^. The rs-fMRI data were collected with 2 mm isotropic resolution, TR = 2s, 53.3 mins (1600 volumes). Details can be seen in Xu et al., 2020. Nineteen macaques with successful preprocessing and surface reconstruction were included in the current study (all males, age = 4.01 ± 0.98 years, weight = 6.61 ±2.04 kilograms).

Macaque data were preprocessed with an HCP-like pipeline ^82^, described elsewhere ^36^. In brief, it included temporal compression, motion correction, 4D global scaling, nuisance regression using white matter (WM), cerebrospinal fluid (CSF), and Friston-24 parameter models, bandpass filtering (0.01 - 0.1 Hz), detrending, and co-registration to the native anatomical space. The data were then projected to the native midcortical surface and smoothed along the surface with FWHM = 3mm. Finally, the preprocessed data were down-sampled to the surface space (with resolution of 10,242 vertices in each hemisphere).

### Parcellations

#### Multimodal parcellation and Cole-Anticevic network

We used multimodal parcellation (MMP) of 360 areas (180 per hemisphere) for humans ^50^. This atlas has been generated using the gradient-based parcellation approach with similar gradient ridges presenting in roughly corresponding locations in both hemispheres, which is suitable for studying asymmetry across homologous parcels. Additionally, based on MMP, we used the Cole-Anticevic Brain-wide Network Partition (CA network), which includes in total 12 functional networks ^51^.

#### ^Desi^kan-Killiany atlas

To ensure our results were reliable we repeated the analysis in humans using a different brain atlas. The Desikan-Killiany atlas ^55^ contains 34 cortical parcels per hemisphere in humans and has high correspondence across two hemispheres.

#### Markov parcellation

For the macaques, we used 91 cortical areas per hemisphere in the Markov M132 architectonic parcellation ^53^. This directed and weighted atlas is generated based on the connectivity profiles. The 91-area parcellation in macaques is valuable for comparison with connectivity analyses in humans.

### Functional connectivity

Cortical time series were averaged within a previously established multi-modal parcellation schemes: for humans the 360-parcelled Glasser atlas (180 per hemisphere) ^50^ and the 91-parcel Markov atlas (91 per hemisphere) for macaques ^53^. To compute the functional connectivity (FC), time-series of cortical parcels were correlated pairwise using the Pearson product moment and then Fisher’s z-transformed in human and macaque data, separately. Individual FC maps were also averaged across four different rs-fMRI sessions for humans ([LR1], [LR2], [RL1], and [RL2]). We computed the FC in four different patterns, both for human and macaque data: FC within the left and right hemispheres (LL intra-hemisphere, RR intra-hemisphere), from the left to right hemisphere (LR inter-hemisphere) and from the right to left hemisphere (RL, inter-hemisphere).

### Connectivity gradients

Next we employed the nonlinear dimensionality reduction technique ^25^ to generate the group level gradients of the mean LL FC across individuals. We then set the group-level gradients as the template and aligned each individual gradient with Procrustes rotation to the template. Finally, the comparative individual functional gradients of each FC pattern were assessed. All steps were accomplished in the Python package Brainspace ^26^.

In brief, the algorithm estimates a low-dimensional embedding from a high-dimensional affinity matrix. Along these low-dimensional axes, or gradients, cortical nodes that are strongly interconnected, by either many suprathreshold edges or few very strong edges, are closer together. Nodes with little connectivity similarly are farther apart. Regions having similar connectivity profiles are embedded together along the gradient axis. The name of this approach, which belongs to the family of graph Laplacians, is derived from the equivalence of the Euclidean distance between points in the diffusion embedded mapping ^25–27^.

The current study selected the first three FC gradients (G1, G2, and G3) that explained 23.3%, 18.1%, and 15.0% of total variance in humans, as well as 18.6%, 14.7%, and 13.0% of total variance in macaques.

### Asymmetry index

To quantify the left and right hemisphere differences, we chose left-right as the asymmetry index (AI) ^4, 9^. In addition, we also calculated the normalized AI with the following formula, (left-right)/(left+right), which is usually used in structural studies to verify whether there is a difference between unnormalized AI and normalized AI. For the intra-hemispheric pattern, the AI was calculated using LL-RR. A positive AI-score meant that the hemispheric feature dominated leftwards, while a negative AI-score dominated rightwards. For the inter- hemispheric pattern we used LR-RL to calculate the AI. Notably, we added ‘minus’ to the AI scores or Cohen’s d scores in the figures in order to conveniently view the lateralization direction.

### Heritability analysis

To map the heritability of functional gradient asymmetry in humans, we used the Sequential Oligogenic Linkage Analysis Routines (SOLAR, v8.5.1b) ^83^. In brief, heritability indicates the impact of genetic relatedness on a phenotype of interest. SOLAR uses maximum likelihood variance decomposition methods to determine the relative importance of familial and environmental influences on a phenotype by modeling the covariance among family members as a function of genetic proximity ^33, 83^. Heritability (i.e. narrow-sense heritability *h*^2^) represents the proportion of the phenotypic variance (σ2p) accounted for by the total additive genetic variance (σ2g), i.e., *h*^2^ = σ2g / σ2p. Phenotypes exhibiting stronger covariances between genetically more similar individuals than between genetically less similar individuals have higher heritability. In this study, we quantified the heritability of asymmetry of functional gradients. We added covariates to our models including age, sex, age^2^, and age × sex.

### Alignment of humans to macaques

To phylogenetically map the asymmetry of functional gradients across macaques and humans, we transformed the human gradients to macaque cortex surface based on a functional joint alignment technique ^36^. This method leverages advances in representing functional organization in high-dimensional common space and provides a transformation between human and macaque cortices, also previously used in ^33, 84^.

In the present study, we aligned Cohen’s d of the human asymmetry index to the macaque surface. Cohens’ d explains the effect size of the asymmetry index. Following the joint alignment, we further computed the Spearman correlation between macaques and humans to evaluate the similarity in asymmetric patterns of the functional gradients. Finally, we compared Cohen’s d between macaques and humans and summarized the results with Markov parcellation ^53^. To illustrate our findings at the functional network level, we projected human networks ^51^ on the macaque surface.

### NeuroSynth meta-analysis

To evaluate the association of function decoding and asymmetry of the principal gradient, we projected the meta-analytical task-based activation along the normalized asymmetry (Cohen’s d) of G1. Our choice for the 24 cognitive domain terms were consistent with ^25^. The activation database we used for meta-analyses was the Neurosynth V3 database ^54^. The surface-based V3 database is available in the github depository (data availability). In the present study, to look at how the right hemisphere and left hemisphere decode functions separately, the leftward normalized asymmetry was put on and the rightward normalized asymmetry was put on the right hemisphere. Other regions became zero. We generated 20 bins along the normalized asymmetry averagely (5% per bin). Thus, each function term had a mean activation z-score per bin. To assess how much the function term was leftward or rightward lateralized, we calculated a weighted score by mean activation (where activation z-score greater than 0.5) multiplied by normalized asymmetry. We roughly regarded this score as the lateralization level. The order of the function terms generated by this calculation reflected the left-right lateralization dominance axis.

## Acknowledgements

We would like to thank the various contributors to the open access databases that our data was downloaded from. Funding: HCP data were provided by the Human Connectome Project, Washington University, the University of Minnesota, and Oxford University Consortium (Principal Investigators: David Van Essen and Kamil Ugurbil; 1U54MH091657) funded by the 16 NIH Institutes and Centers that support the NIH Blueprint for Neuroscience Research; and by the McDonnell Center for Systems Neuroscience at Washington University. Additional personnel support provided by the Center for the Developing Brain at the Child Mind Institute, as well as NIMH R01MH081218, R01MH083246, and R21MH084126. Project support is also provided by the NKI Center for Advanced Brain Imaging (CABI), the Brain Research Foundation (Chicago, IL), and the Stavros Niarchos Foundation. This study was supported by the Deutsche Forschungsgemeinschaft (DFG, EI 816/21-1), the National Institute of Mental Health (R01-MH074457), the Helmholtz Portfolio Theme “Supercomputing and Modeling for the Human Brain” and the European Union’s Horizon 2020 Research and Innovation Program under Grant Agreement No. 785907 (HBP SGA2). S.L.V was supported by Max Planck Gesellschaft (Otto Hahn award). B.C.B acknowledges support from the SickKids Foundation (NI17-039), the National Sciences and Engineering Research Council of Canada (NSERC; Discovery-1304413), CIHR (FDN154298), Azrieli Center for Autism Research (ACAR), an MNI-Cambridge collaboration grant, and the Canada Research Chairs program. Last, this work was funded in part by Helmholtz Association’s Initiative and Networking Fund under the Helmholtz International Lab grant agreement InterLabs-0015, and the Canada First Research Excellence Fund (CFREF Competition 2, 2015-2016) awarded to the Healthy Brains, Healthy Lives initiative at McGill University, through the Helmholtz International BigBrain Analytics and Learning Laboratory (HIBALL), including S.L.V. and B.C.B.. B.W. was supported by the International Max Planck Research School on Neuroscience of Communication: Function, Structure, and Plasticity (IMPRS NeuroCom).

## Author contributions

B.W. and S.L.V. conceived and designed the analysis, performed the analysis, wrote the draft manuscript and revised the manuscript. Ş.B. aided in data analysis. T.X. provided fMRI data of macaques and gave critical comments. R.A.I.B provided fMRI data of the UK Biobank. Ş.B., T.X., H.L.S., R.A.I.B. and B.C.B. helped writing and revising the manuscript.

## Competing interests

All co-authors declare no conflict of interests.

## Materials & correspondence

This article is corresponding to Bin Wan (Ph.D. student) and Dr. Sofie Valk (Ph.D.) at Cognitive Neurogenetics (CNG) lab, Max Planck Institute for Human Cognitive and Brain Sciences, Leipzig, Germany, and Research Center Jülich, Jülich, Germany.

## Data availability

All human data analyzed in this manuscript were obtained from the open-access HCP young adult sample (HCP; www.humanconnectome.org/), UK Biobank (UKB, https://www.ukbiobank.ac.uk/). Macaque data came from PRIME-DE (http://fcon_1000.projects.nitrc.org/indi/indiPRIME.html). Gradient analyses and visualization were performed using the Python package Brainspace (https://brainspace.readthedocs.io/en/latest/index.html, ^26^. Heritability analyses were performed using Solar Eclipse 8.5.1b (www.solar-eclipse-genetics.org). Task-based function association analyses were based on NeuroSynth (https://neurosynth.org/, ^54^. Full statistical scripts can be found at https://github.com/CNG-=LAB/cngopen/tree/main/asymmetry_functional_gradients/scripts.

## Supplementary Results

### Robustness analyses

Complementing our main AI calculation (L-R), we additionally used AI_norm (L-R)/(L+R), with rescaling the distribution of gradients to positive values, to explore whether our results were robust with respect to AI calculation (**Supplementary Fig. S6**). We found that for G1, asymmetric effects were highly correlated with the main asymmetric effects (Spearman *r* _intra- hemisphere_ = 0.9998, *P* < 0.001; *r* _inter-hemisphere_ = 0.999, *P* _uncorrected_ < 0.001). High correlation was also found in G2 (Spearman *r* _intra-hemisphere_ = 0.9990, *P* _uncorrected_ < 0.001; *r* _inter-hemisphere_ = 0.9993, *P* _uncorrected_ < 0.001) and in G3 (Spearman *r* _intra-hemisphere_ = 0.9997, *P* _uncorrected_ < 0.001; *r* _inter- hemisphere_ = 0.9997, *P* _uncorrected_ < 0.001). The spatial map of mean AI and *P* value with FDR correction was also found similar to that of mean AI_norm and *P* value with FDR correction.

To test the robustness of our findings with respect to the parcellation approach, we employed Desikan-Killiany atlas ^1^ to generate the asymmetry of functional gradients. This is a symmetric atlas containing 34 parcels per hemisphere. Overall, for the intra-hemispheric pattern G1 showed similar hemispheric patterns as observed in our main results when using the Desikan-Killiany atlas. In particular, the posterior cluster between middle and superior temporal gyrus and Broca’s area showed leftward asymmetry, whereas dorso-lateral prefrontal regions showed rightward asymmetry (**Supplementary Fig. S7**). However, we observed more details are shown in multi-modal parcellation. Similar patterns were observed in inter-hemispheric asymmetry of functional organization when using the Desikan-Killiany atlas.

We also used an additional sample (UK Biobank, UKB) to verify whether asymmetry of functional organization is present in other samples. We included 34,604 subjects’ imaging data in UKB with good quality. After computing Cohen’s d of asymmetric effects in UKB, to account for differences in sample size, we performed a group level correlation between HCP with UKB (**Supplementary Fig. S8**). We observed a high correlation between the LL functional gradient between HCP and UKB (Spearman *r* _G1_ = -0.591, *P* _uncorrected_ < 0.001; Spearman *r* _G2_ = 0.312, *P* _uncorrected_ < 0.001; Spearman *r* _G3_ = 0.771, *P* _uncorrected_ < 0.001). Thus, we flipped UKB LL G1 direction to make it more consistent with HCP (now *r* _G1_ = 0.591, *P* _uncorrected_ < 0.001). For the G1 of the intra-hemispheric FC pattern, we observed a correlation with our findings in the HCP sample (Spearman *r* = 0.269, *P* _gen_ < 0.022). However, association between HCP and UKB was not significant for inter-hemispheric asymmetry patterns (Spearman *r* = 0.043, *P* _gen_ = 0.852). All the networks showed significant asymmetry in UKB. However, we found that language and default mode networks showed leftward asymmetry (as in HCP), but the frontoparietal network did not show rightward asymmetry.

---

**Fig. S1.**
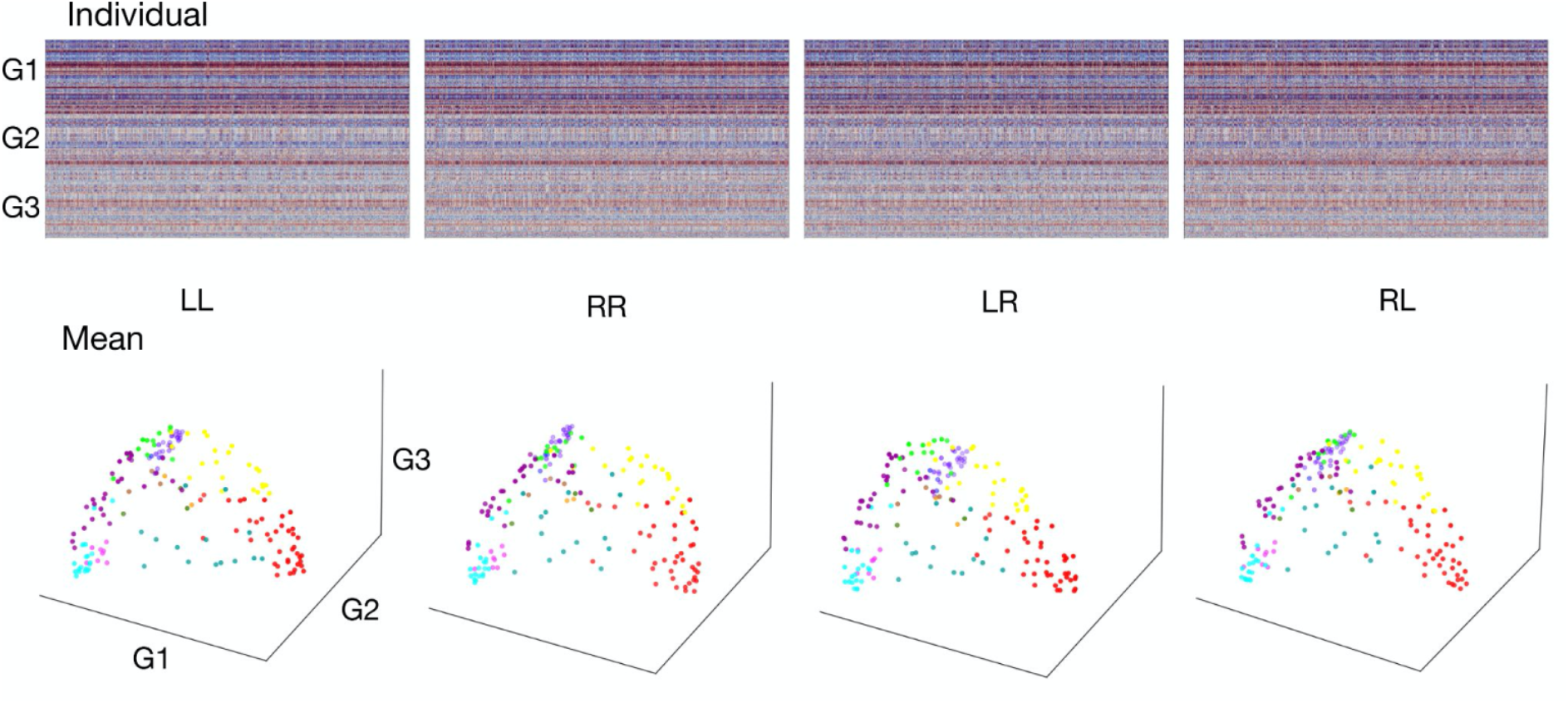
Individual gradients of each FC pattern.

**Fig. S2.**
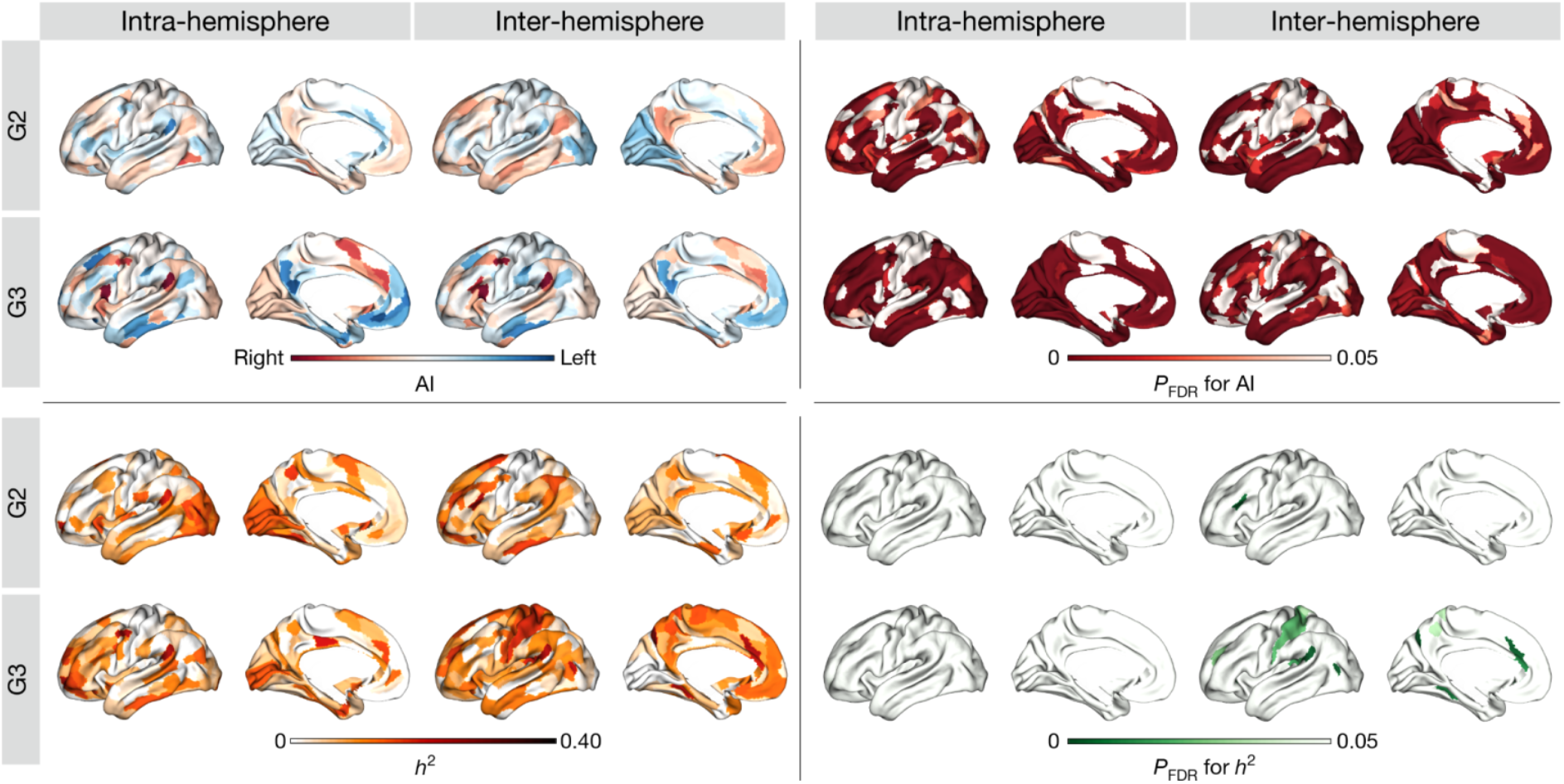
Mean asymmetry index (AI), heritability (*h*^2^), and *P*_FDR_ of G2 and G3.

**Fig. S3.**
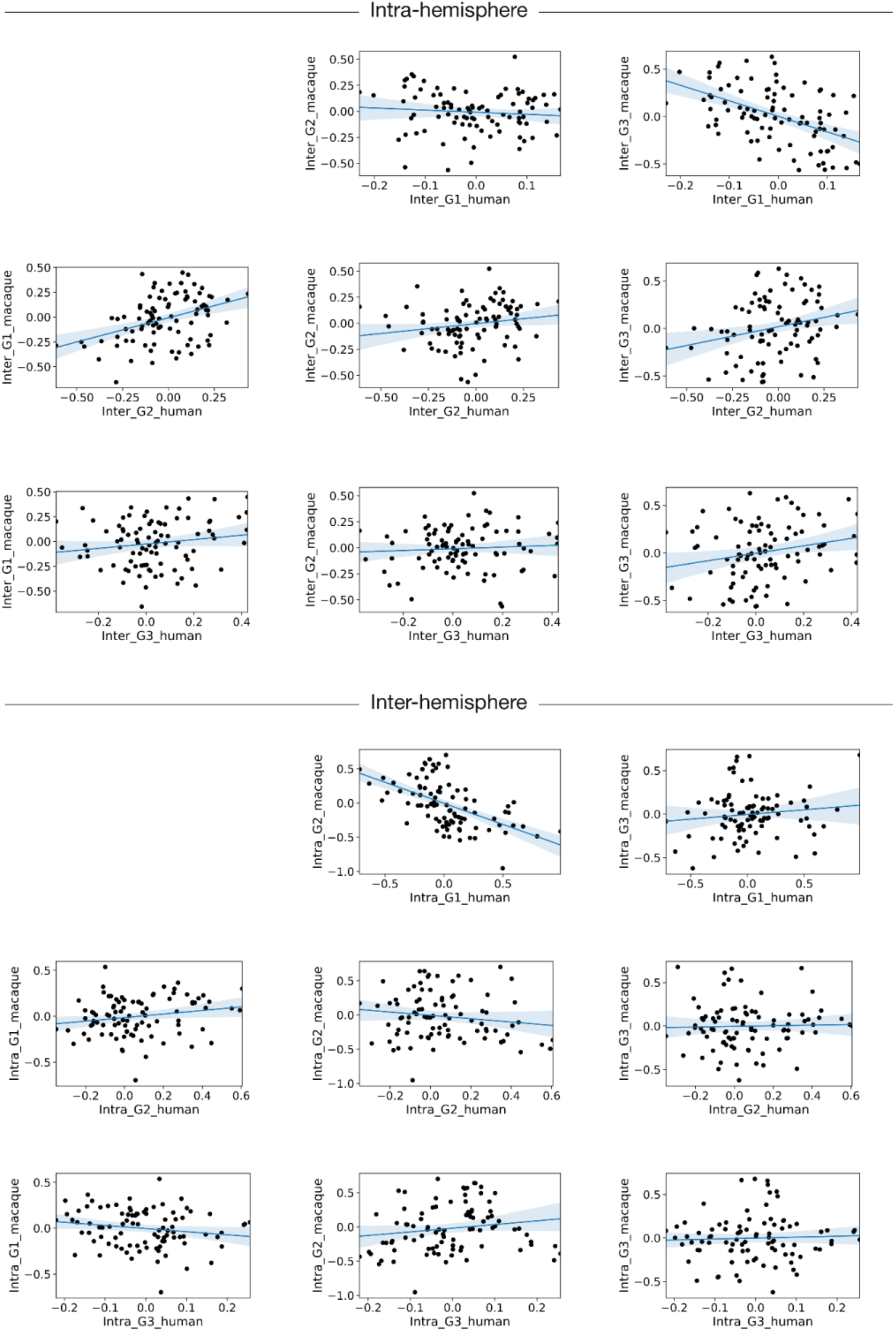
Similarity of asymmetry of functional gradients in macaques and humans.

**Fig. S4.**
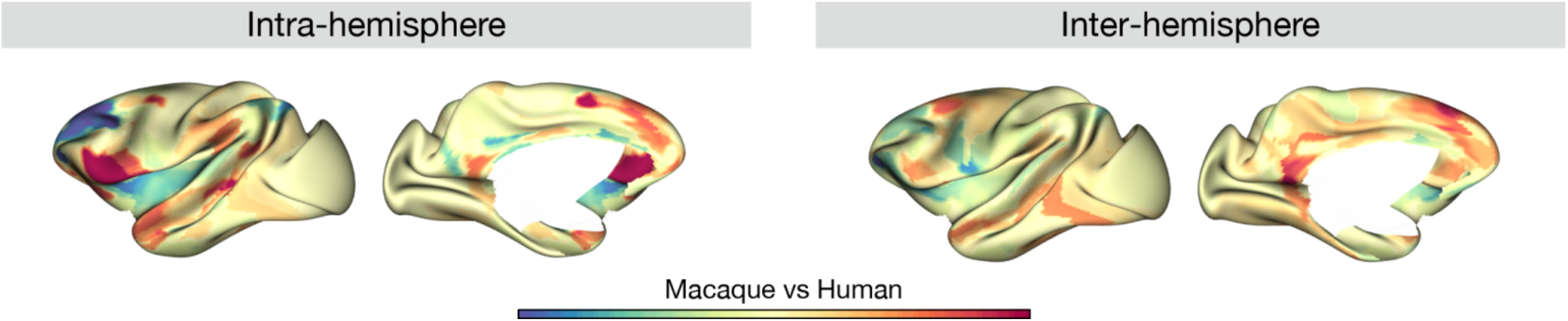
Macaque versus human region-wise difference maps based on the normalized (Cohen’s d) asymmetry of G1.

**Fig. S5.**
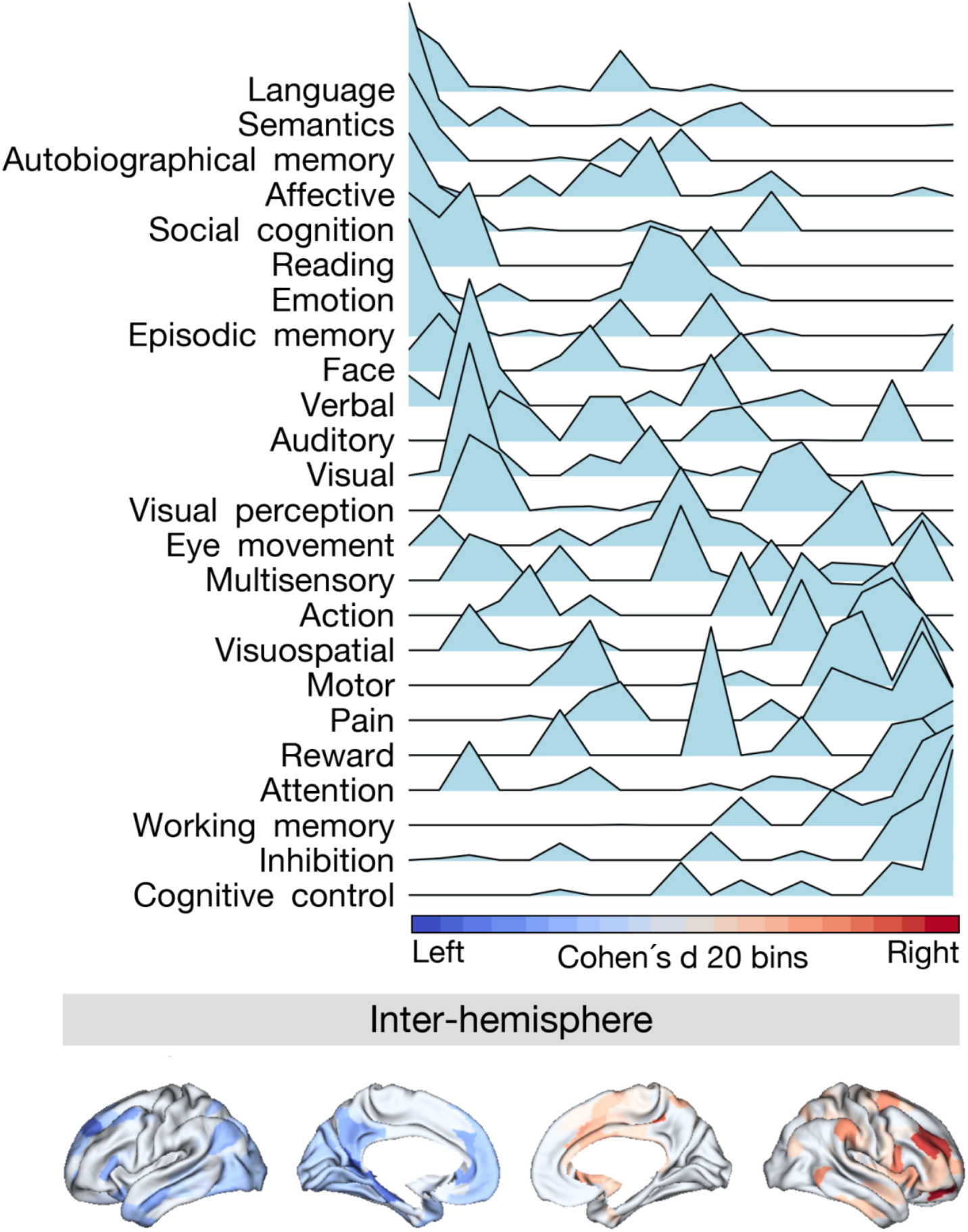
Projection of meta-analytical task-based function along normalized (Cohen’s d) asymmetry of G1 (inter-hemisphere).

**Fig. S6.**
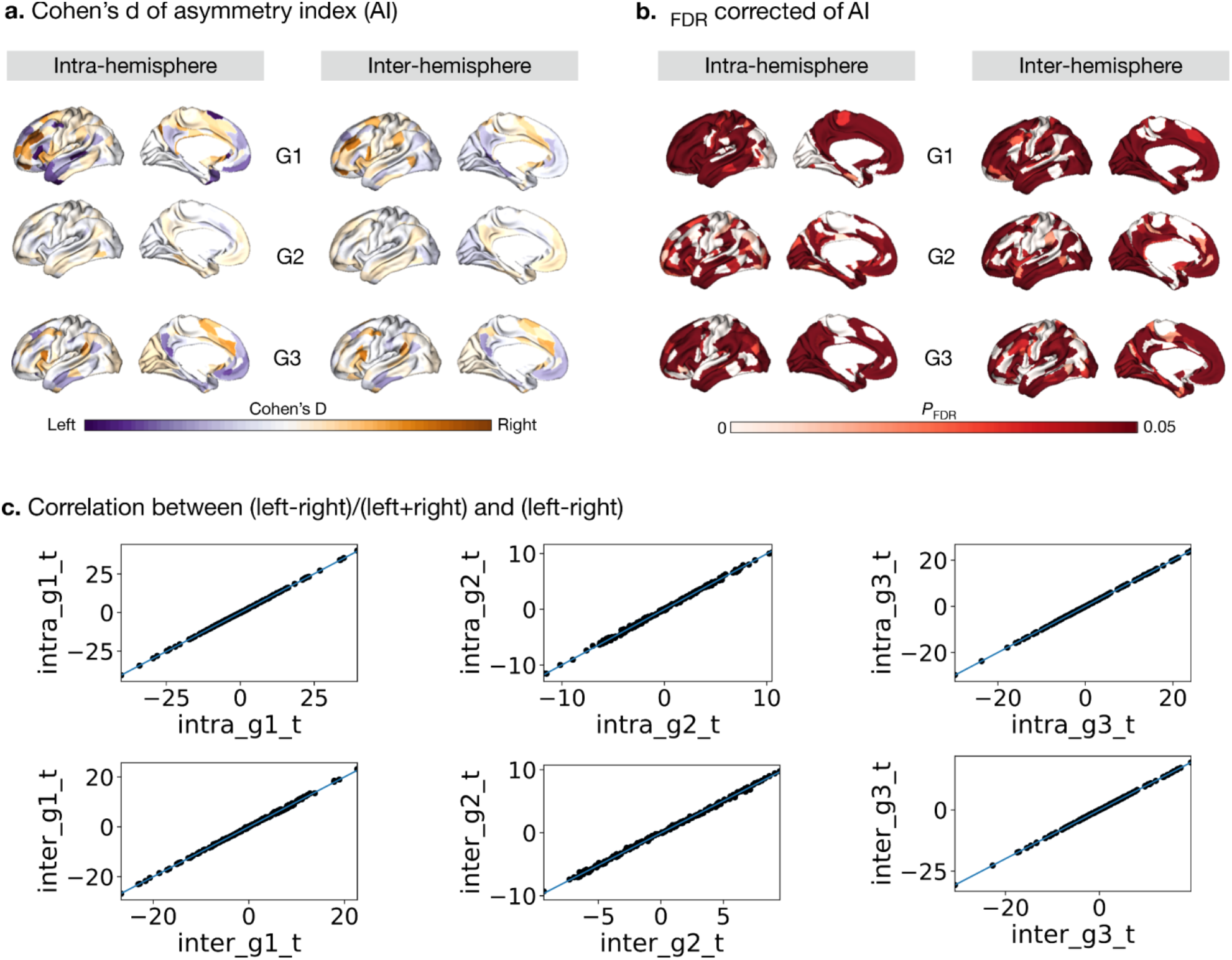
Cohen’s d (yellow-purple) and *P*_FDR_ (red) of asymmetry index with (left- right)/(left+right).

**Fig. S7.**
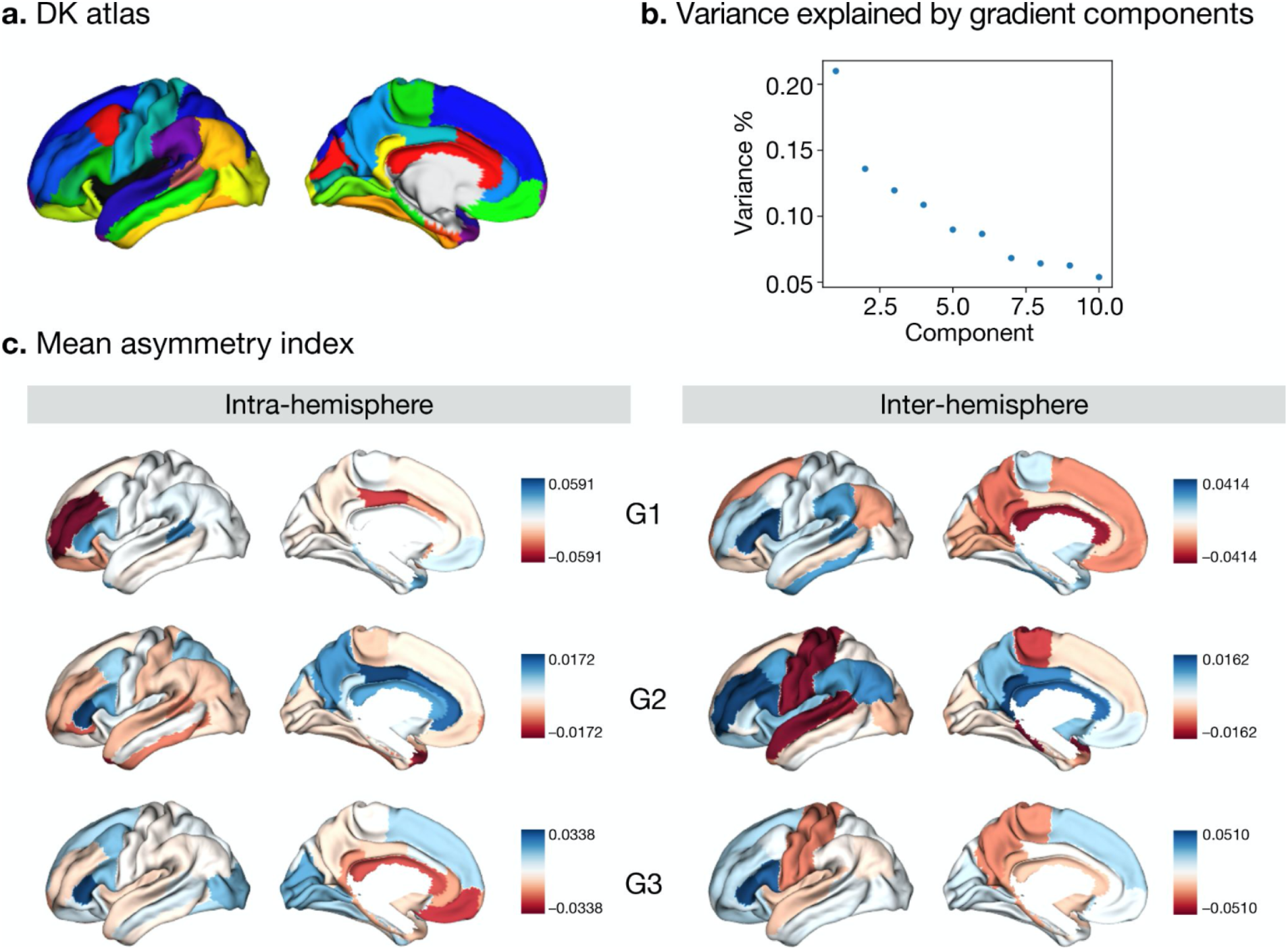
Asymmetry using Desikan-Killiany (DK) atlas^1^.

**Fig. S8.**
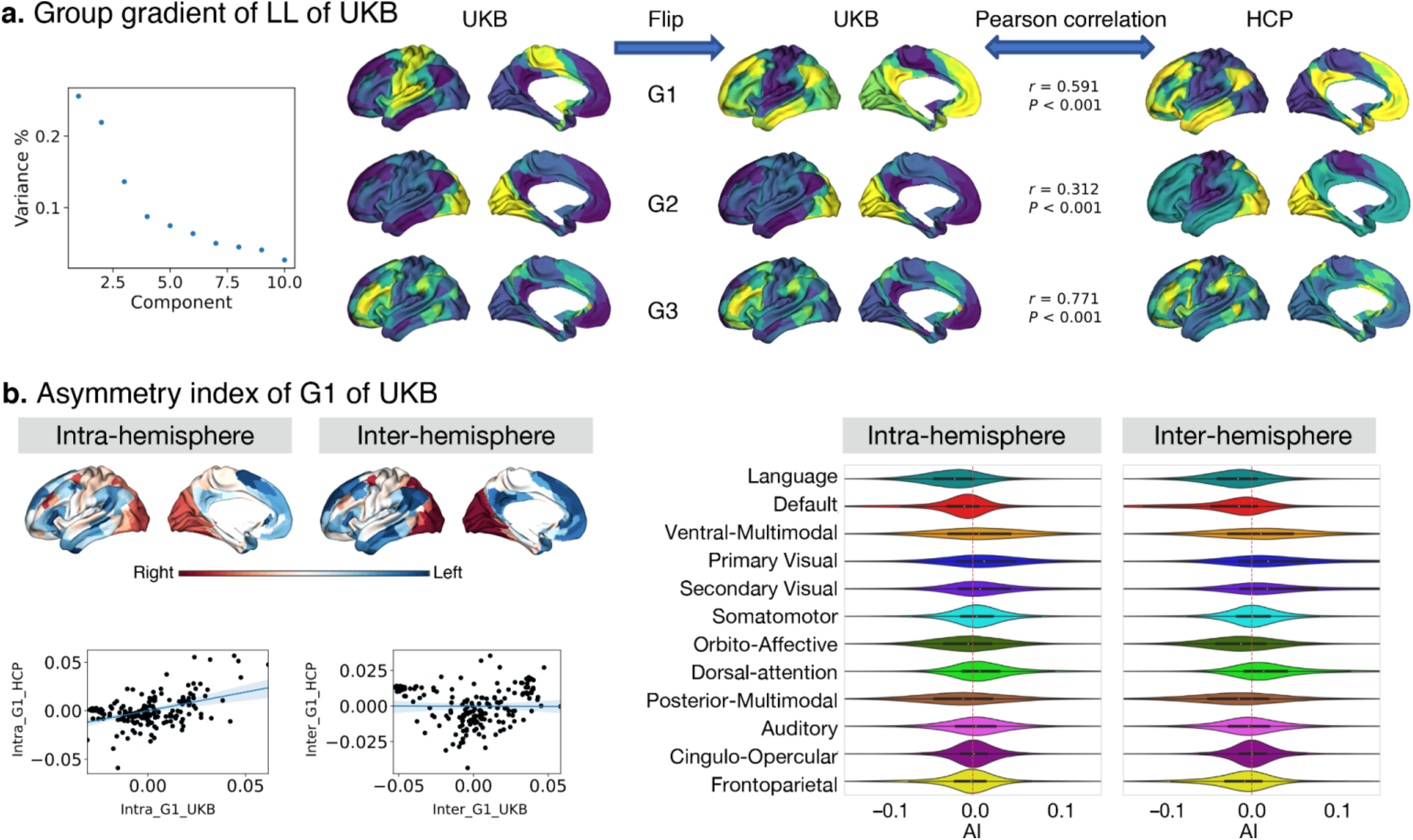
Asymmetry using UK Biobank sample (n = 34,604).

**Table S1.**
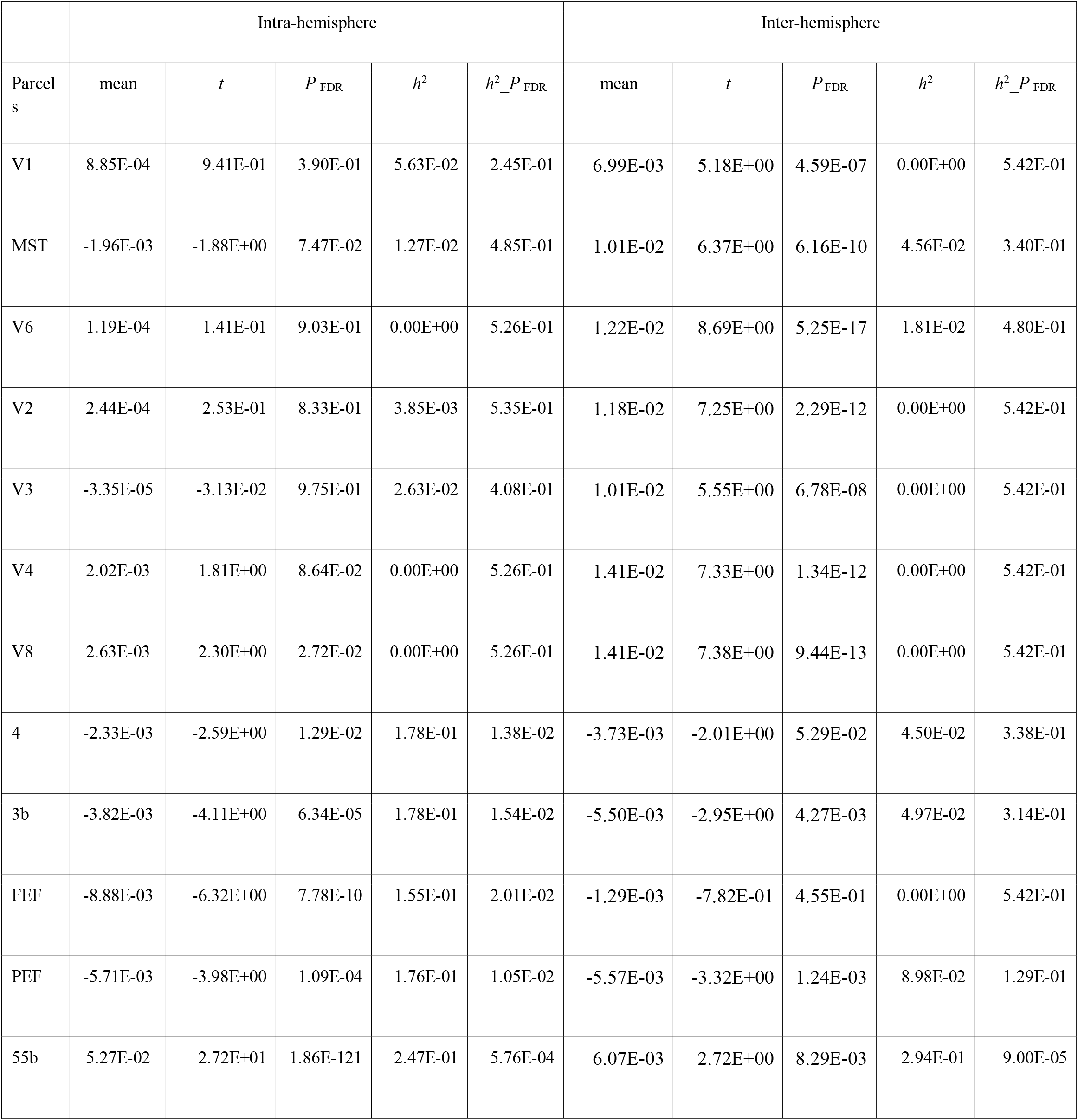
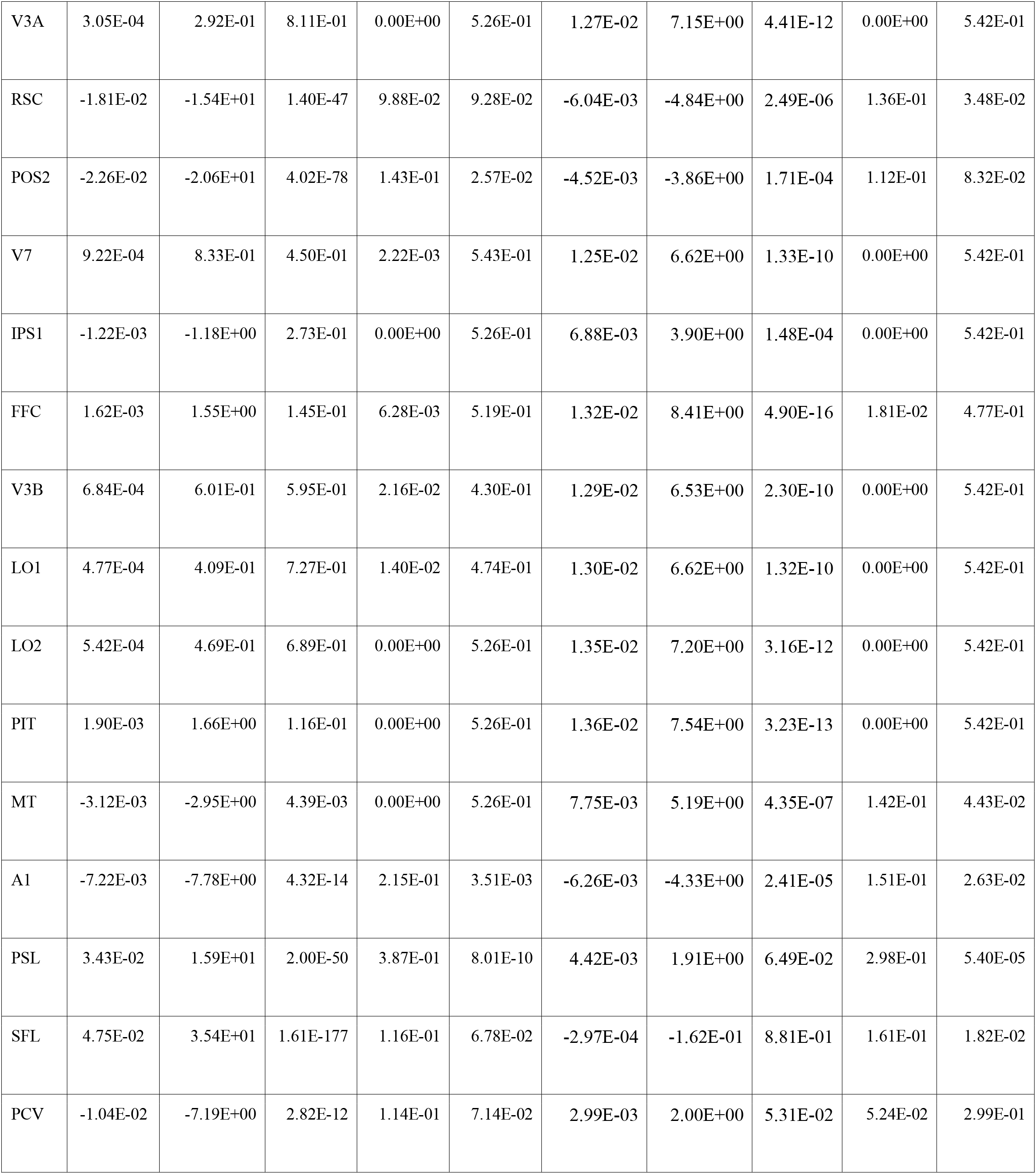
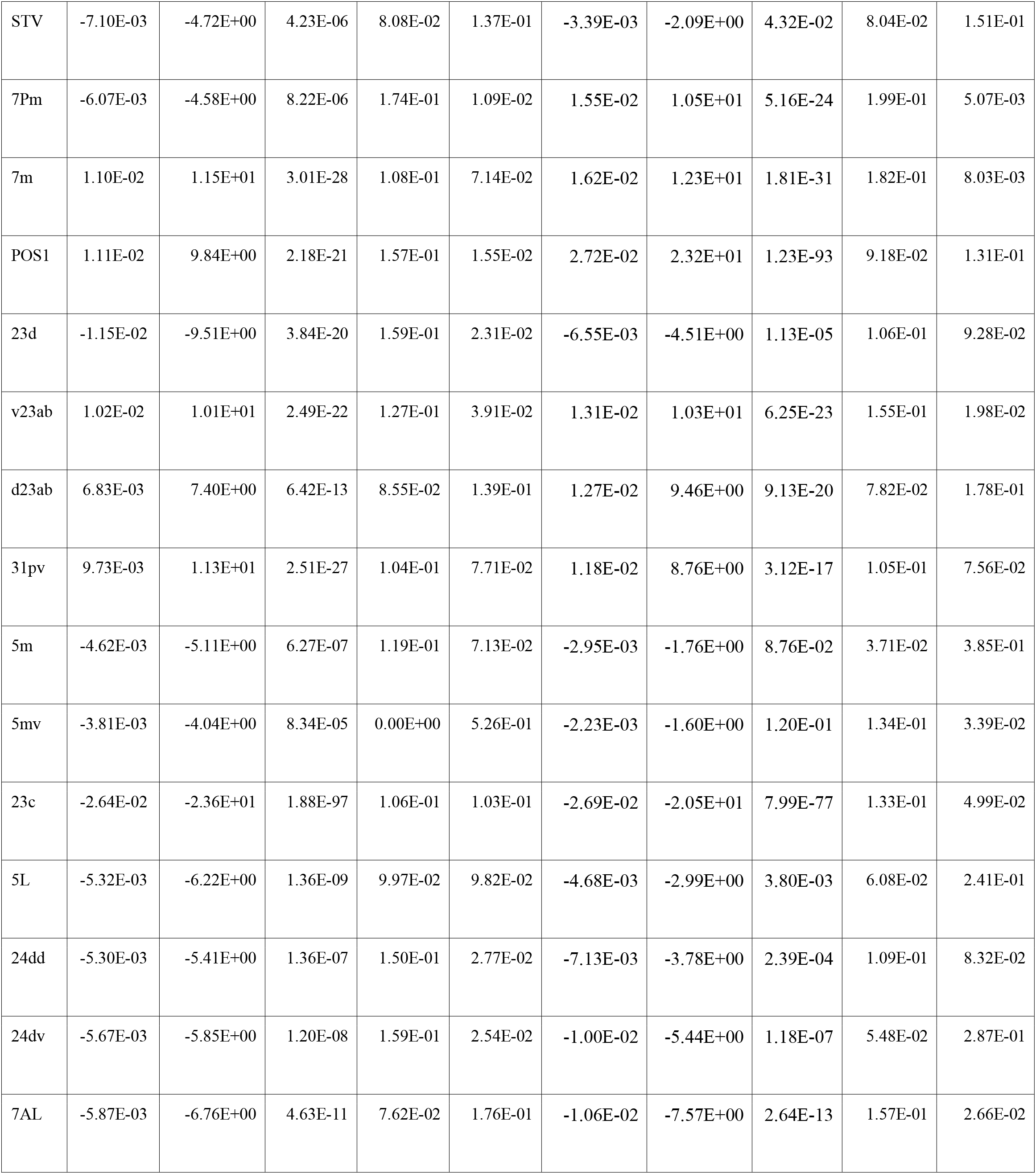
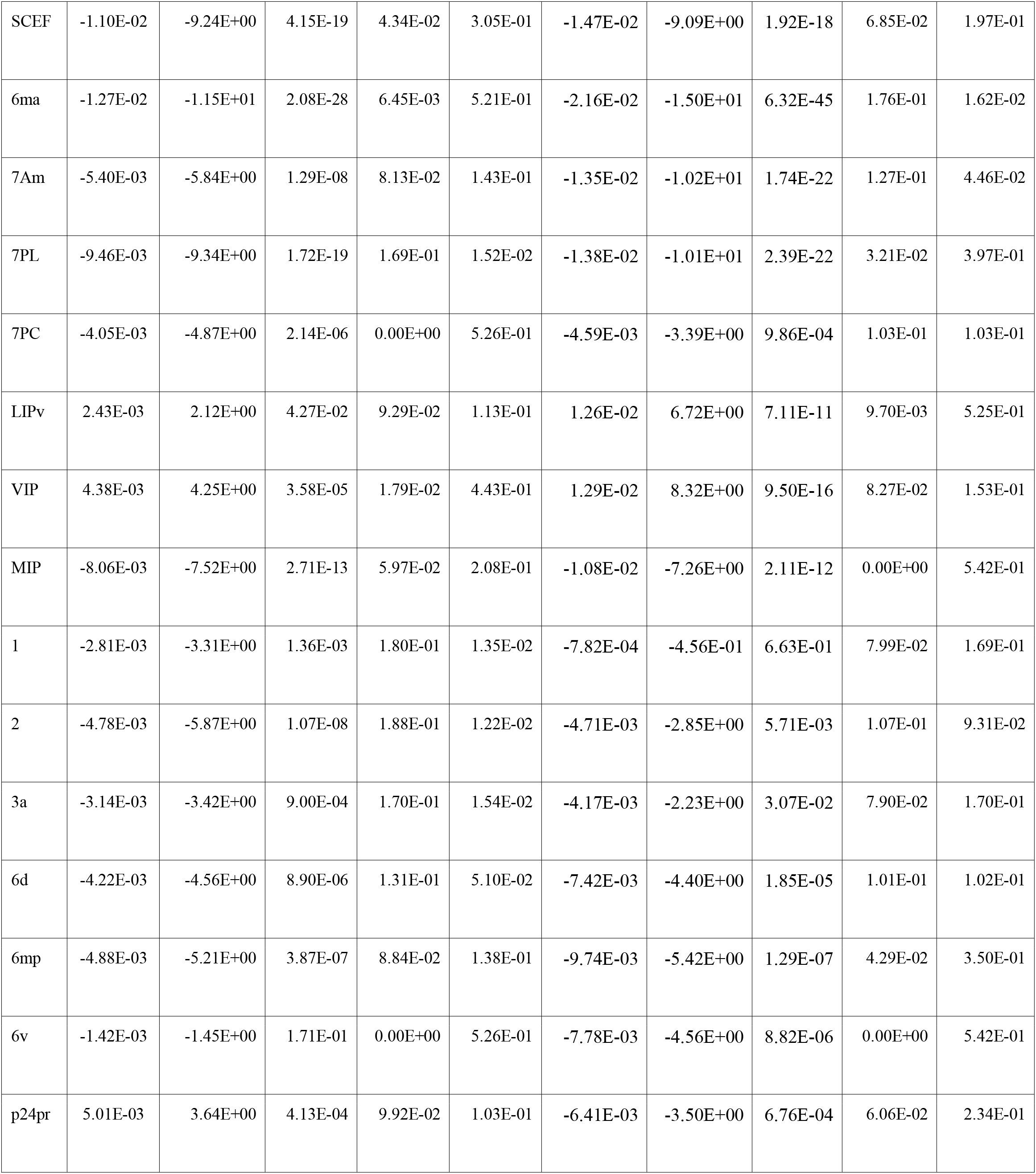
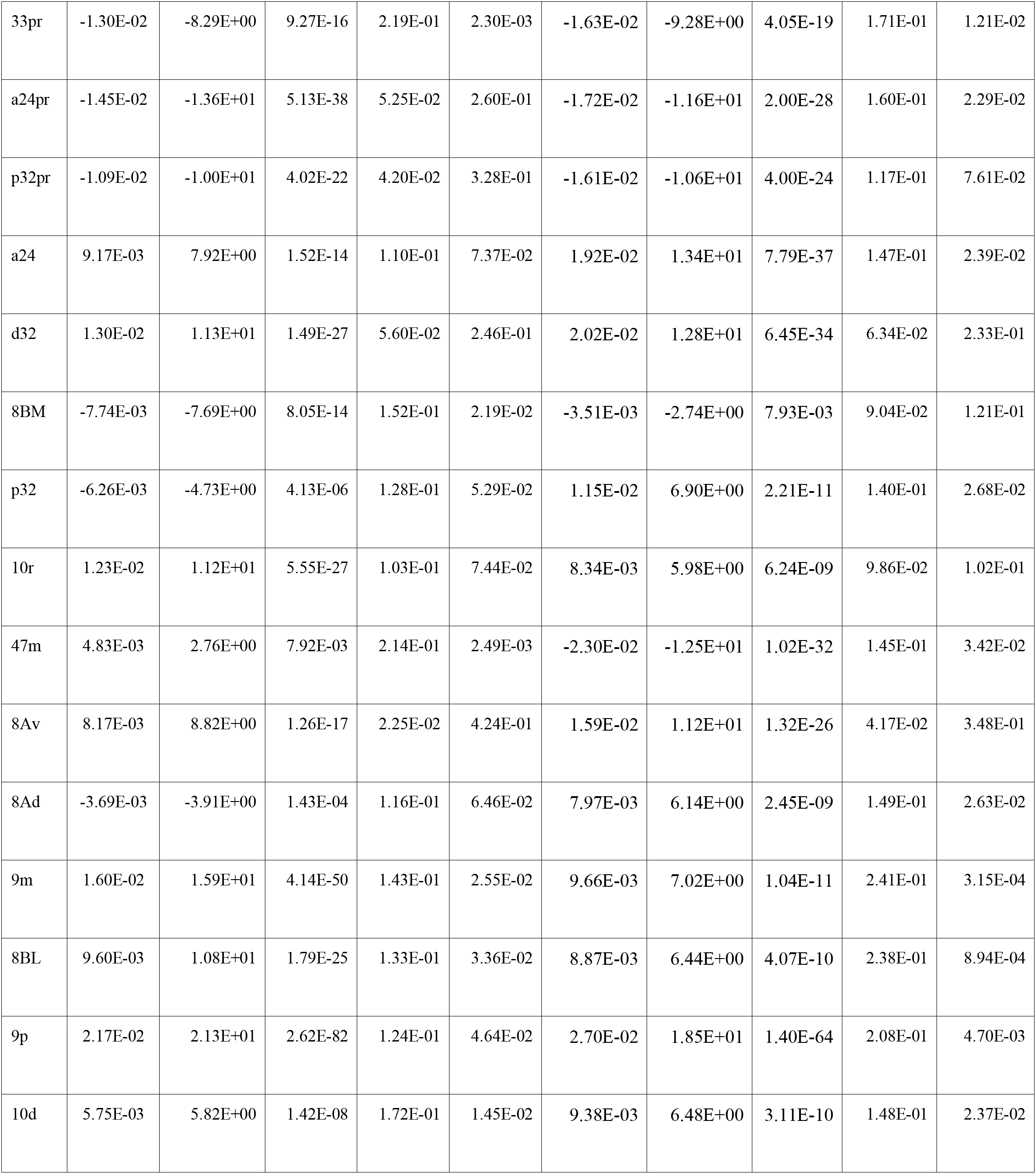
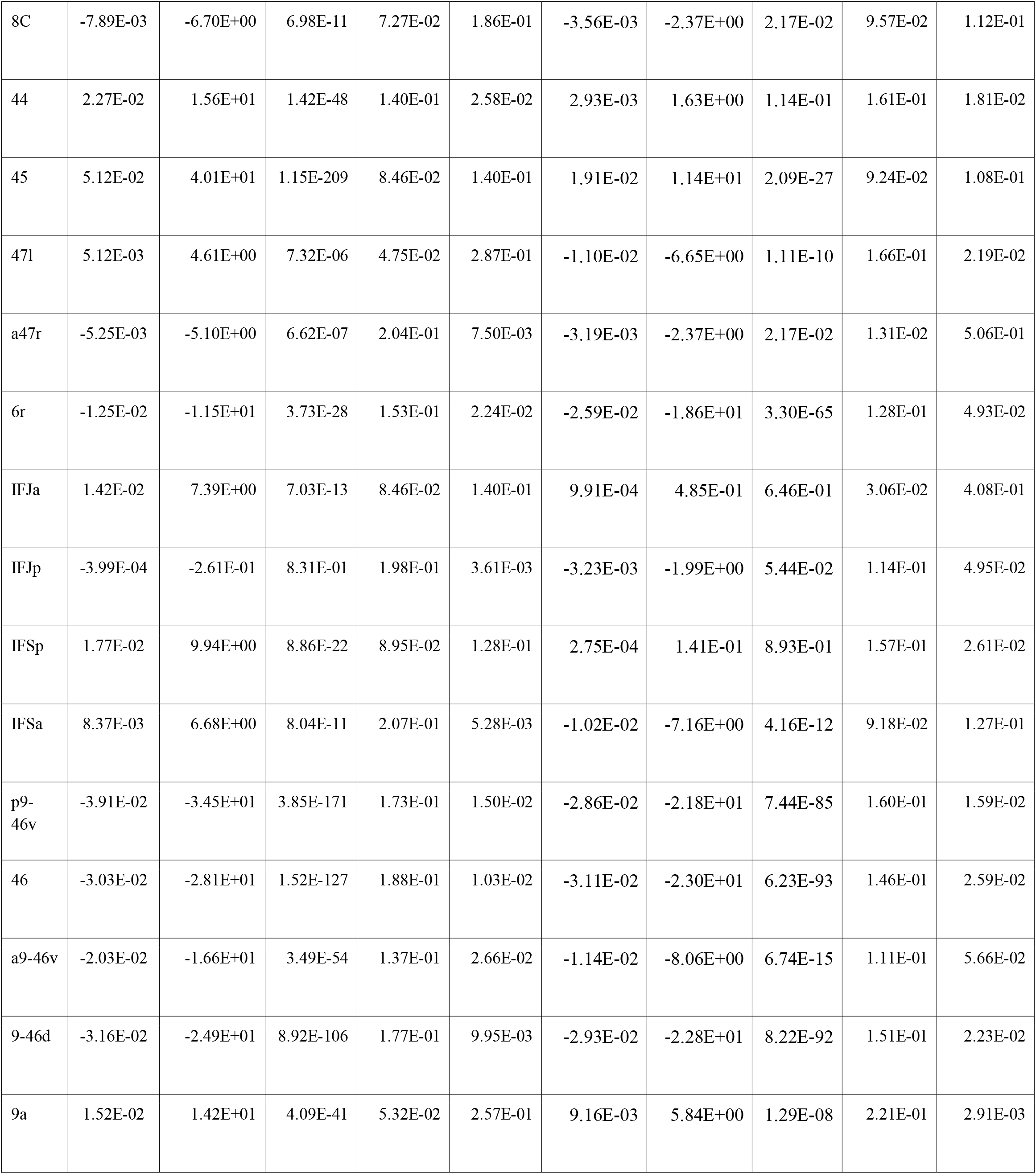
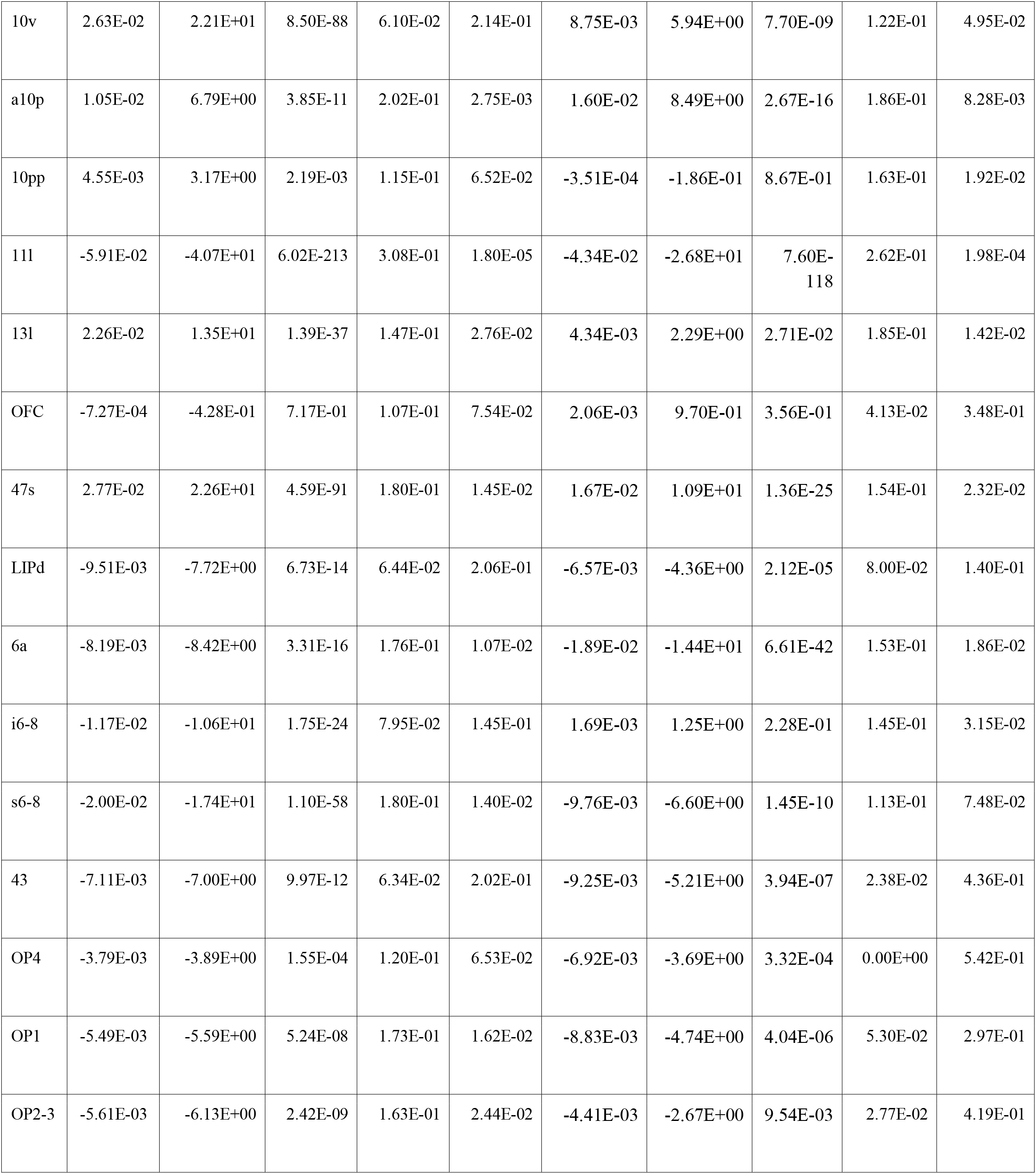
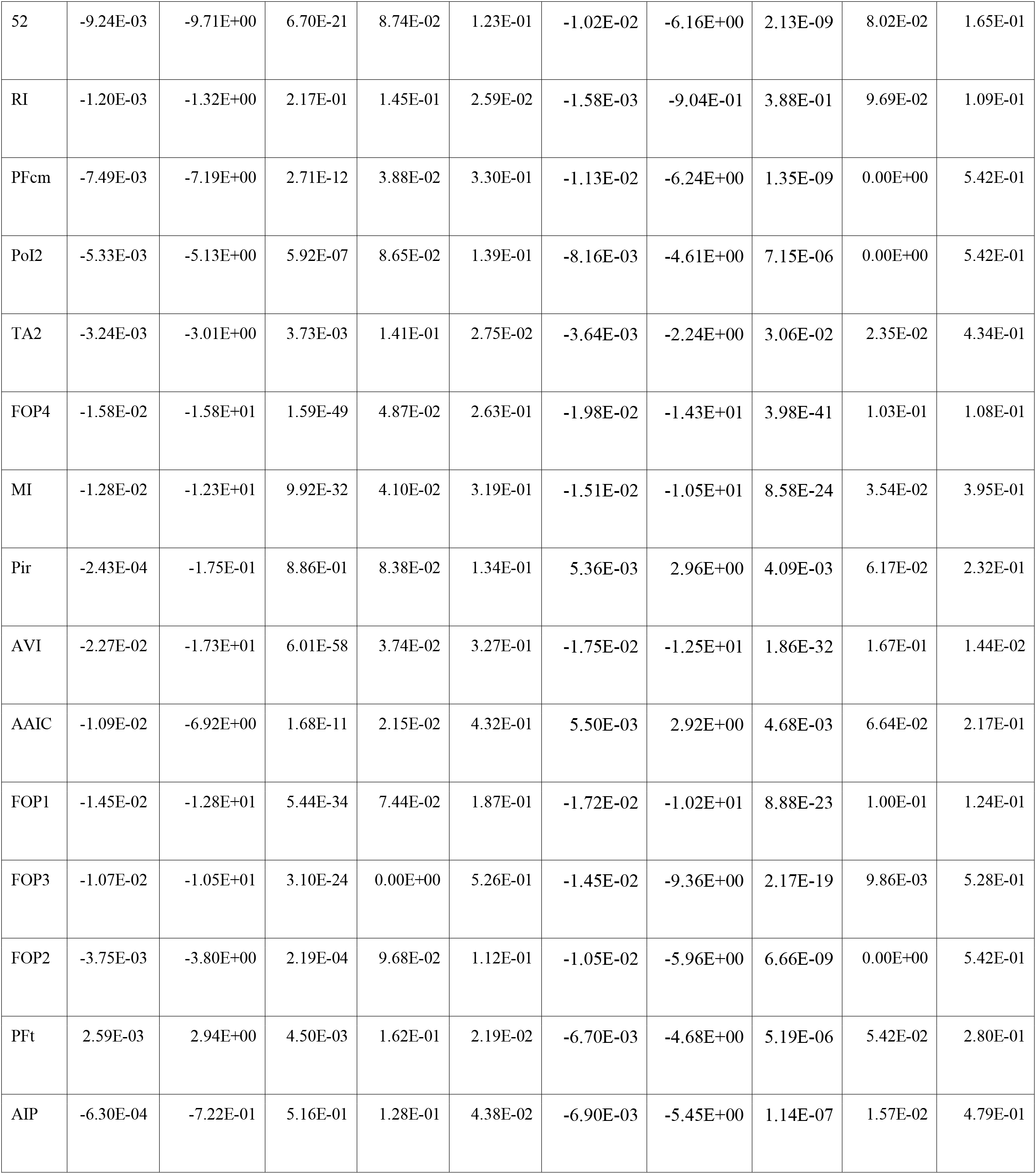
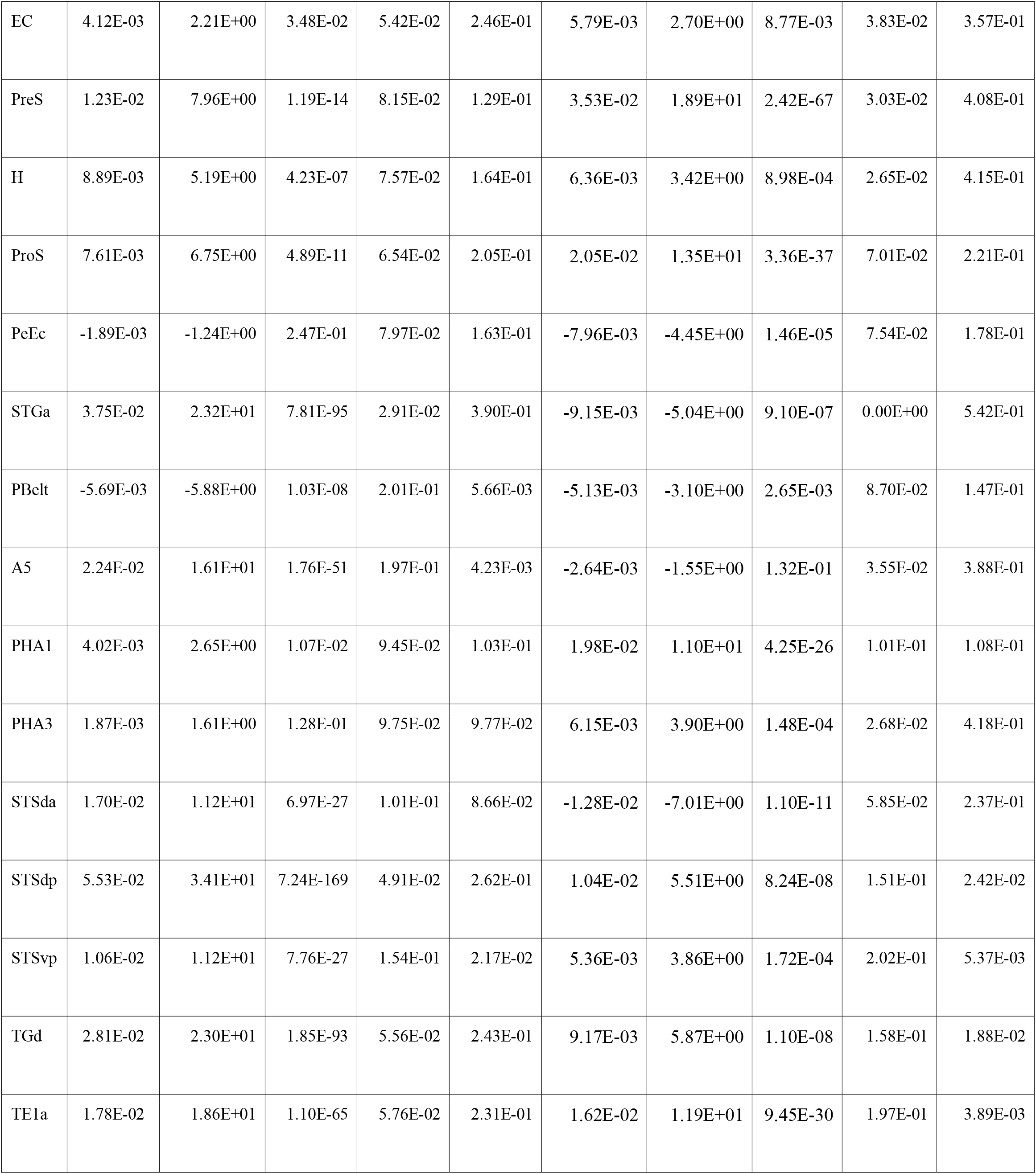
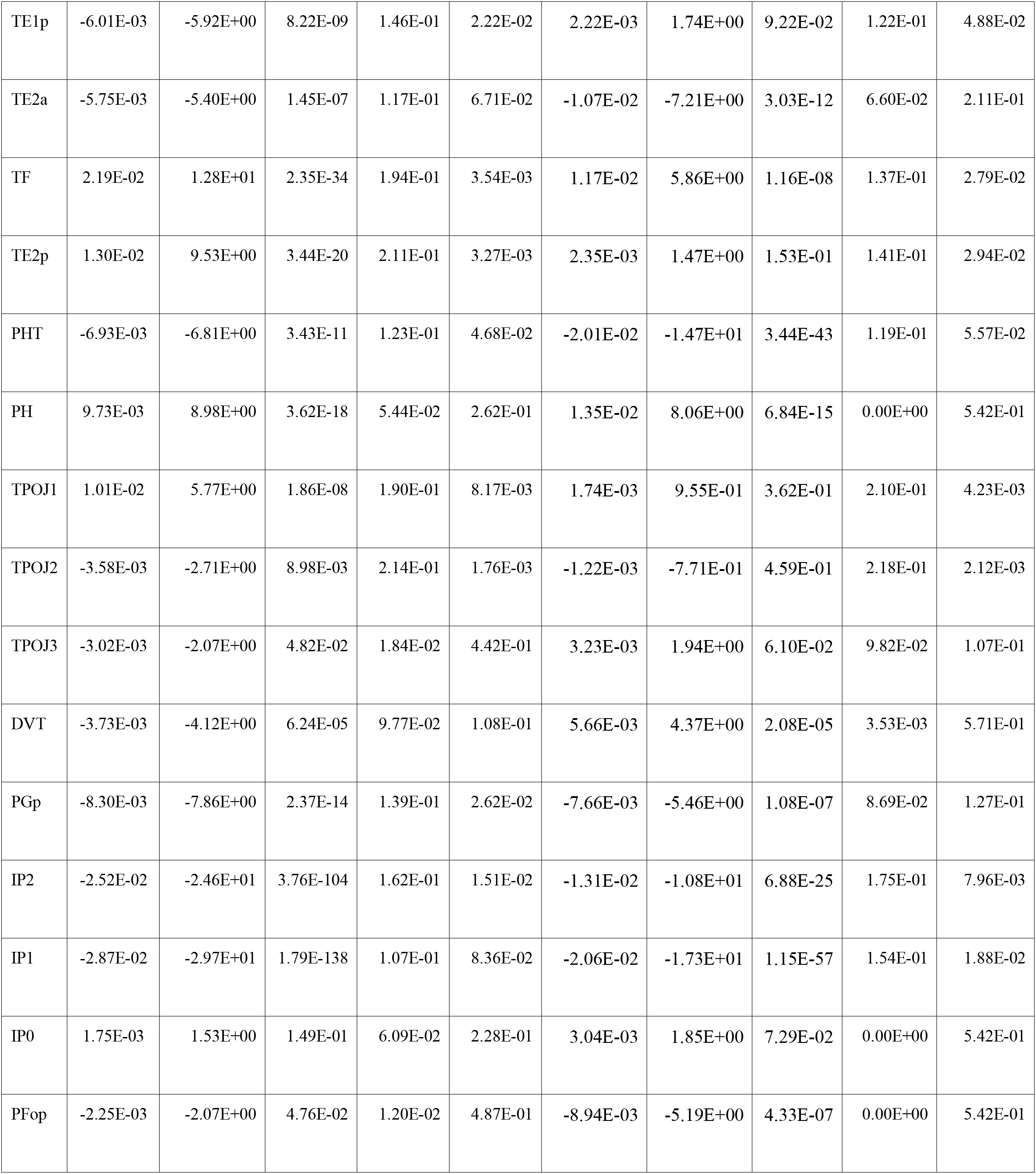
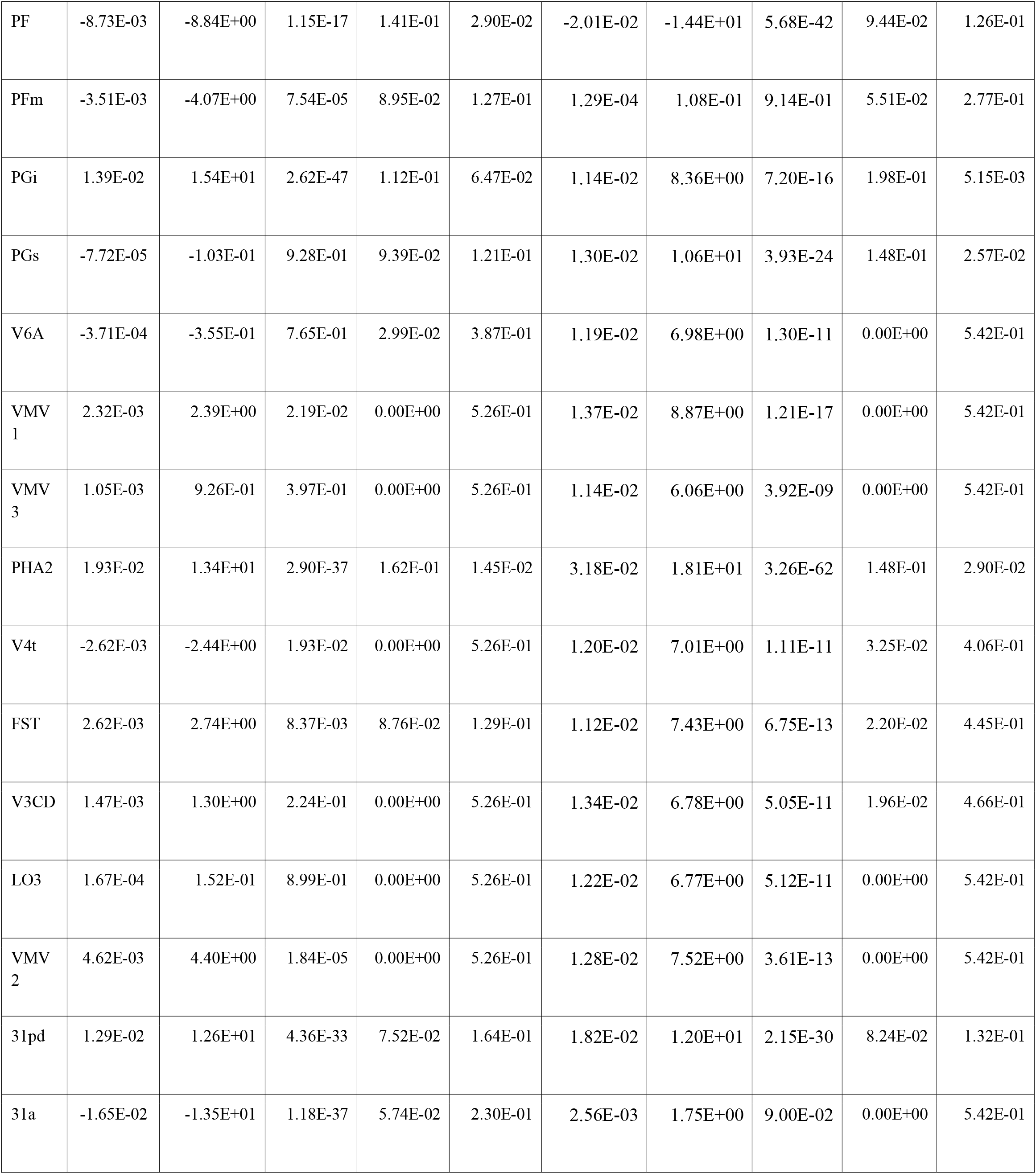
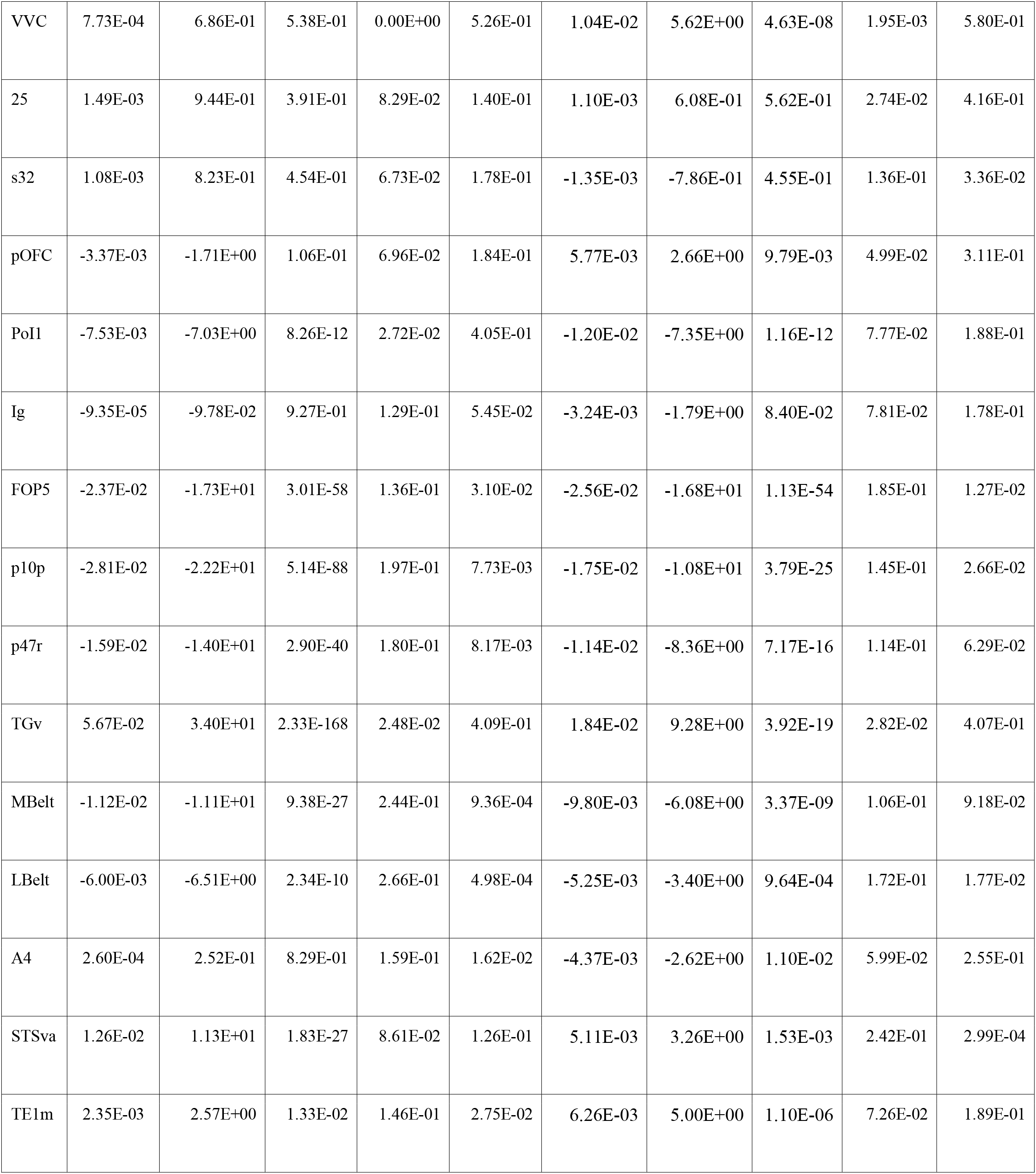
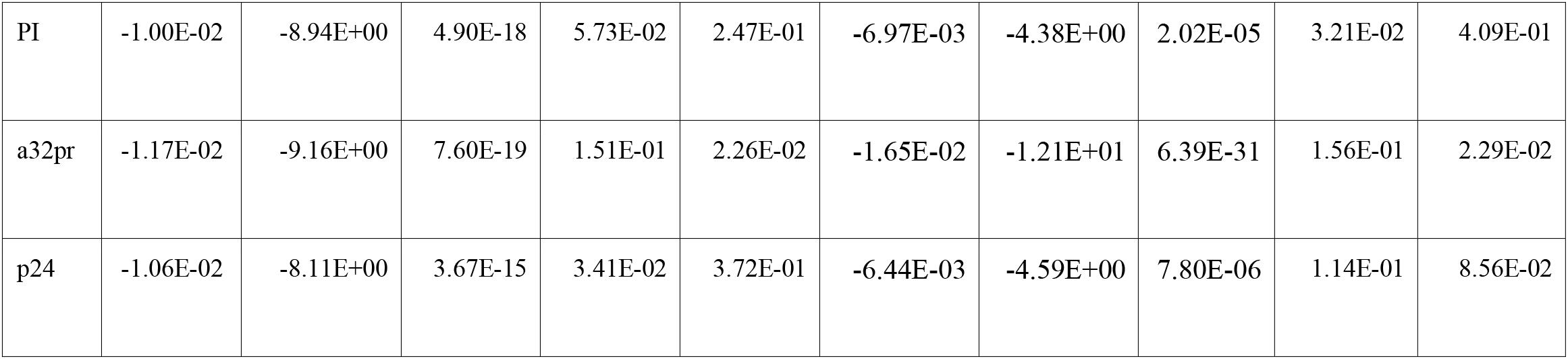
Summary of asymmetry index and heritability of G1, for detailed description of parcel names, please see Glasser et al., 2016^2.^

|  | Intra-hemisphere |  |  |  |  | Inter-hemisphere |  |  |  |  |
| --- | --- | --- | --- | --- | --- | --- | --- | --- | --- | --- |
| Parcel<br>s | mean | $t$ | $P_{\text{FDR}}$ | $h^2$ | $h^2_{\text{P}_{\text{FDR}}}$ | mean | $t$ | $P_{\text{FDR}}$ | $h^2$ | $h^2_{\text{P}_{\text{FDR}}}$ |
| V1 | 8.85E-04 | 9.41E-01 | 3.90E-01 | 5.63E-02 | 2.45E-01 | 6.99E-03 | 5.18E+00 | 4.59E-07 | 0.00E+00 | 5.42E-01 |
| MST | -1.96E-03 | -1.88E+00 | 7.47E-02 | 1.27E-02 | 4.85E-01 | 1.01E-02 | 6.37E+00 | 6.16E-10 | 4.56E-02 | 3.40E-01 |
| V6 | 1.19E-04 | 1.41E-01 | 9.03E-01 | 0.00E+00 | 5.26E-01 | 1.22E-02 | 8.69E+00 | 5.25E-17 | 1.81E-02 | 4.80E-01 |
| V2 | 2.44E-04 | 2.53E-01 | 8.33E-01 | 3.85E-03 | 5.35E-01 | 1.18E-02 | 7.25E+00 | 2.29E-12 | 0.00E+00 | 5.42E-01 |
| V3 | -3.35E-05 | -3.13E-02 | 9.75E-01 | 2.63E-02 | 4.08E-01 | 1.01E-02 | 5.55E+00 | 6.78E-08 | 0.00E+00 | 5.42E-01 |
| V4 | 2.02E-03 | 1.81E+00 | 8.64E-02 | 0.00E+00 | 5.26E-01 | 1.41E-02 | 7.33E+00 | 1.34E-12 | 0.00E+00 | 5.42E-01 |
| V8 | 2.63E-03 | 2.30E+00 | 2.72E-02 | 0.00E+00 | 5.26E-01 | 1.41E-02 | 7.38E+00 | 9.44E-13 | 0.00E+00 | 5.42E-01 |
| 4 | -2.33E-03 | -2.59E+00 | 1.29E-02 | 1.78E-01 | 1.38E-02 | -3.73E-03 | -2.01E+00 | 5.29E-02 | 4.50E-02 | 3.38E-01 |
| 3b | -3.82E-03 | -4.11E+00 | 6.34E-05 | 1.78E-01 | 1.54E-02 | -5.50E-03 | -2.95E+00 | 4.27E-03 | 4.97E-02 | 3.14E-01 |
| FEF | -8.88E-03 | -6.32E+00 | 7.78E-10 | 1.55E-01 | 2.01E-02 | -1.29E-03 | -7.82E-01 | 4.55E-01 | 0.00E+00 | 5.42E-01 |
| PEF | -5.71E-03 | -3.98E+00 | 1.09E-04 | 1.76E-01 | 1.05E-02 | -5.57E-03 | -3.32E+00 | 1.24E-03 | 8.98E-02 | 1.29E-01 |
| 55b | 5.27E-02 | 2.72E+01 | 1.86E-121 | 2.47E-01 | 5.76E-04 | 6.07E-03 | 2.72E+00 | 8.29E-03 | 2.94E-01 | 9.00E-05 |
| V3A | 3.05E-04 | 2.92E-01 | 8.11E-01 | 0.00E+00 | 5.26E-01 | 1.27E-02 | 7.15E+00 | 4.41E-12 | 0.00E+00 | 5.42E-01 |
| RSC | -1.81E-02 | -1.54E+01 | 1.40E-47 | 9.88E-02 | 9.28E-02 | -6.04E-03 | -4.84E+00 | 2.49E-06 | 1.36E-01 | 3.48E-02 |
| POS2 | -2.26E-02 | -2.06E+01 | 4.02E-78 | 1.43E-01 | 2.57E-02 | -4.52E-03 | -3.86E+00 | 1.71E-04 | 1.12E-01 | 8.32E-02 |
| V7 | 9.22E-04 | 8.33E-01 | 4.50E-01 | 2.22E-03 | 5.43E-01 | 1.25E-02 | 6.62E+00 | 1.33E-10 | 0.00E+00 | 5.42E-01 |
| IPS1 | -1.22E-03 | -1.18E+00 | 2.73E-01 | 0.00E+00 | 5.26E-01 | 6.88E-03 | 3.90E+00 | 1.48E-04 | 0.00E+00 | 5.42E-01 |
| FFC | 1.62E-03 | 1.55E+00 | 1.45E-01 | 6.28E-03 | 5.19E-01 | 1.32E-02 | 8.41E+00 | 4.90E-16 | 1.81E-02 | 4.77E-01 |
| V3B | 6.84E-04 | 6.01E-01 | 5.95E-01 | 2.16E-02 | 4.30E-01 | 1.29E-02 | 6.53E+00 | 2.30E-10 | 0.00E+00 | 5.42E-01 |
| LO1 | 4.77E-04 | 4.09E-01 | 7.27E-01 | 1.40E-02 | 4.74E-01 | 1.30E-02 | 6.62E+00 | 1.32E-10 | 0.00E+00 | 5.42E-01 |
| LO2 | 5.42E-04 | 4.69E-01 | 6.89E-01 | 0.00E+00 | 5.26E-01 | 1.35E-02 | 7.20E+00 | 3.16E-12 | 0.00E+00 | 5.42E-01 |
| PIT | 1.90E-03 | 1.66E+00 | 1.16E-01 | 0.00E+00 | 5.26E-01 | 1.36E-02 | 7.54E+00 | 3.23E-13 | 0.00E+00 | 5.42E-01 |
| MT | -3.12E-03 | -2.95E+00 | 4.39E-03 | 0.00E+00 | 5.26E-01 | 7.75E-03 | 5.19E+00 | 4.35E-07 | 1.42E-01 | 4.43E-02 |
| A1 | -7.22E-03 | -7.78E+00 | 4.32E-14 | 2.15E-01 | 3.51E-03 | -6.26E-03 | -4.33E+00 | 2.41E-05 | 1.51E-01 | 2.63E-02 |
| PSL | 3.43E-02 | 1.59E+01 | 2.00E-50 | 3.87E-01 | 8.01E-10 | 4.42E-03 | 1.91E+00 | 6.49E-02 | 2.98E-01 | 5.40E-05 |
| SFL | 4.75E-02 | 3.54E+01 | 1.61E-177 | 1.16E-01 | 6.78E-02 | -2.97E-04 | -1.62E-01 | 8.81E-01 | 1.61E-01 | 1.82E-02 |
| PCV | -1.04E-02 | -7.19E+00 | 2.82E-12 | 1.14E-01 | 7.14E-02 | 2.99E-03 | 2.00E+00 | 5.31E-02 | 5.24E-02 | 2.99E-01 |
| STV | -7.10E-03 | -4.72E+00 | 4.23E-06 | 8.08E-02 | 1.37E-01 | -3.39E-03 | -2.09E+00 | 4.32E-02 | 8.04E-02 | 1.51E-01 |
| 7Pm | -6.07E-03 | -4.58E+00 | 8.22E-06 | 1.74E-01 | 1.09E-02 | 1.55E-02 | 1.05E+01 | 5.16E-24 | 1.99E-01 | 5.07E-03 |
| 7m | 1.10E-02 | 1.15E+01 | 3.01E-28 | 1.08E-01 | 7.14E-02 | 1.62E-02 | 1.23E+01 | 1.81E-31 | 1.82E-01 | 8.03E-03 |
| POS1 | 1.11E-02 | 9.84E+00 | 2.18E-21 | 1.57E-01 | 1.55E-02 | 2.72E-02 | 2.32E+01 | 1.23E-93 | 9.18E-02 | 1.31E-01 |
| 23d | -1.15E-02 | -9.51E+00 | 3.84E-20 | 1.59E-01 | 2.31E-02 | -6.55E-03 | -4.51E+00 | 1.13E-05 | 1.06E-01 | 9.28E-02 |
| v23ab | 1.02E-02 | 1.01E+01 | 2.49E-22 | 1.27E-01 | 3.91E-02 | 1.31E-02 | 1.03E+01 | 6.25E-23 | 1.55E-01 | 1.98E-02 |
| d23ab | 6.83E-03 | 7.40E+00 | 6.42E-13 | 8.55E-02 | 1.39E-01 | 1.27E-02 | 9.46E+00 | 9.13E-20 | 7.82E-02 | 1.78E-01 |
| 31pv | 9.73E-03 | 1.13E+01 | 2.51E-27 | 1.04E-01 | 7.71E-02 | 1.18E-02 | 8.76E+00 | 3.12E-17 | 1.05E-01 | 7.56E-02 |
| 5m | -4.62E-03 | -5.11E+00 | 6.27E-07 | 1.19E-01 | 7.13E-02 | -2.95E-03 | -1.76E+00 | 8.76E-02 | 3.71E-02 | 3.85E-01 |
| 5mv | -3.81E-03 | -4.04E+00 | 8.34E-05 | 0.00E+00 | 5.26E-01 | -2.23E-03 | -1.60E+00 | 1.20E-01 | 1.34E-01 | 3.39E-02 |
| 23c | -2.64E-02 | -2.36E+01 | 1.88E-97 | 1.06E-01 | 1.03E-01 | -2.69E-02 | -2.05E+01 | 7.99E-77 | 1.33E-01 | 4.99E-02 |
| 5L | -5.32E-03 | -6.22E+00 | 1.36E-09 | 9.97E-02 | 9.82E-02 | -4.68E-03 | -2.99E+00 | 3.80E-03 | 6.08E-02 | 2.41E-01 |
| 24dd | -5.30E-03 | -5.41E+00 | 1.36E-07 | 1.50E-01 | 2.77E-02 | -7.13E-03 | -3.78E+00 | 2.39E-04 | 1.09E-01 | 8.32E-02 |
| 24dv | -5.67E-03 | -5.85E+00 | 1.20E-08 | 1.59E-01 | 2.54E-02 | -1.00E-02 | -5.44E+00 | 1.18E-07 | 5.48E-02 | 2.87E-01 |
| 7AL | -5.87E-03 | -6.76E+00 | 4.63E-11 | 7.62E-02 | 1.76E-01 | -1.06E-02 | -7.57E+00 | 2.64E-13 | 1.57E-01 | 2.66E-02 |
| SCEF | -1.10E-02 | -9.24E+00 | 4.15E-19 | 4.34E-02 | 3.05E-01 | -1.47E-02 | -9.09E+00 | 1.92E-18 | 6.85E-02 | 1.97E-01 |
| 6ma | -1.27E-02 | -1.15E+01 | 2.08E-28 | 6.45E-03 | 5.21E-01 | -2.16E-02 | -1.50E+01 | 6.32E-45 | 1.76E-01 | 1.62E-02 |
| 7Am | -5.40E-03 | -5.84E+00 | 1.29E-08 | 8.13E-02 | 1.43E-01 | -1.35E-02 | -1.02E+01 | 1.74E-22 | 1.27E-01 | 4.46E-02 |
| 7PL | -9.46E-03 | -9.34E+00 | 1.72E-19 | 1.69E-01 | 1.52E-02 | -1.38E-02 | -1.01E+01 | 2.39E-22 | 3.21E-02 | 3.97E-01 |
| 7PC | -4.05E-03 | -4.87E+00 | 2.14E-06 | 0.00E+00 | 5.26E-01 | -4.59E-03 | -3.39E+00 | 9.86E-04 | 1.03E-01 | 1.03E-01 |
| LIPv | 2.43E-03 | 2.12E+00 | 4.27E-02 | 9.29E-02 | 1.13E-01 | 1.26E-02 | 6.72E+00 | 7.11E-11 | 9.70E-03 | 5.25E-01 |
| VIP | 4.38E-03 | 4.25E+00 | 3.58E-05 | 1.79E-02 | 4.43E-01 | 1.29E-02 | 8.32E+00 | 9.50E-16 | 8.27E-02 | 1.53E-01 |
| MIP | -8.06E-03 | -7.52E+00 | 2.71E-13 | 5.97E-02 | 2.08E-01 | -1.08E-02 | -7.26E+00 | 2.11E-12 | 0.00E+00 | 5.42E-01 |
| 1 | -2.81E-03 | -3.31E+00 | 1.36E-03 | 1.80E-01 | 1.35E-02 | -7.82E-04 | -4.56E-01 | 6.63E-01 | 7.99E-02 | 1.69E-01 |
| 2 | -4.78E-03 | -5.87E+00 | 1.07E-08 | 1.88E-01 | 1.22E-02 | -4.71E-03 | -2.85E+00 | 5.71E-03 | 1.07E-01 | 9.31E-02 |
| 3a | -3.14E-03 | -3.42E+00 | 9.00E-04 | 1.70E-01 | 1.54E-02 | -4.17E-03 | -2.23E+00 | 3.07E-02 | 7.90E-02 | 1.70E-01 |
| 6d | -4.22E-03 | -4.56E+00 | 8.90E-06 | 1.31E-01 | 5.10E-02 | -7.42E-03 | -4.40E+00 | 1.85E-05 | 1.01E-01 | 1.02E-01 |
| 6mp | -4.88E-03 | -5.21E+00 | 3.87E-07 | 8.84E-02 | 1.38E-01 | -9.74E-03 | -5.42E+00 | 1.29E-07 | 4.29E-02 | 3.50E-01 |
| 6v | -1.42E-03 | -1.45E+00 | 1.71E-01 | 0.00E+00 | 5.26E-01 | -7.78E-03 | -4.56E+00 | 8.82E-06 | 0.00E+00 | 5.42E-01 |
| p24pr | 5.01E-03 | 3.64E+00 | 4.13E-04 | 9.92E-02 | 1.03E-01 | -6.41E-03 | -3.50E+00 | 6.76E-04 | 6.06E-02 | 2.34E-01 |
| 33pr | -1.30E-02 | -8.29E+00 | 9.27E-16 | 2.19E-01 | 2.30E-03 | -1.63E-02 | -9.28E+00 | 4.05E-19 | 1.71E-01 | 1.21E-02 |
| a24pr | -1.45E-02 | -1.36E+01 | 5.13E-38 | 5.25E-02 | 2.60E-01 | -1.72E-02 | -1.16E+01 | 2.00E-28 | 1.60E-01 | 2.29E-02 |
| p32pr | -1.09E-02 | -1.00E+01 | 4.02E-22 | 4.20E-02 | 3.28E-01 | -1.61E-02 | -1.06E+01 | 4.00E-24 | 1.17E-01 | 7.61E-02 |
| a24 | 9.17E-03 | 7.92E+00 | 1.52E-14 | 1.10E-01 | 7.37E-02 | 1.92E-02 | 1.34E+01 | 7.79E-37 | 1.47E-01 | 2.39E-02 |
| d32 | 1.30E-02 | 1.13E+01 | 1.49E-27 | 5.60E-02 | 2.46E-01 | 2.02E-02 | 1.28E+01 | 6.45E-34 | 6.34E-02 | 2.33E-01 |
| 8BM | -7.74E-03 | -7.69E+00 | 8.05E-14 | 1.52E-01 | 2.19E-02 | -3.51E-03 | -2.74E+00 | 7.93E-03 | 9.04E-02 | 1.21E-01 |
| p32 | -6.26E-03 | -4.73E+00 | 4.13E-06 | 1.28E-01 | 5.29E-02 | 1.15E-02 | 6.90E+00 | 2.21E-11 | 1.40E-01 | 2.68E-02 |
| 10r | 1.23E-02 | 1.12E+01 | 5.55E-27 | 1.03E-01 | 7.44E-02 | 8.34E-03 | 5.98E+00 | 6.24E-09 | 9.86E-02 | 1.02E-01 |
| 47m | 4.83E-03 | 2.76E+00 | 7.92E-03 | 2.14E-01 | 2.49E-03 | -2.30E-02 | -1.25E+01 | 1.02E-32 | 1.45E-01 | 3.42E-02 |
| 8Av | 8.17E-03 | 8.82E+00 | 1.26E-17 | 2.25E-02 | 4.24E-01 | 1.59E-02 | 1.12E+01 | 1.32E-26 | 4.17E-02 | 3.48E-01 |
| 8Ad | -3.69E-03 | -3.91E+00 | 1.43E-04 | 1.16E-01 | 6.46E-02 | 7.97E-03 | 6.14E+00 | 2.45E-09 | 1.49E-01 | 2.63E-02 |
| 9m | 1.60E-02 | 1.59E+01 | 4.14E-50 | 1.43E-01 | 2.55E-02 | 9.66E-03 | 7.02E+00 | 1.04E-11 | 2.41E-01 | 3.15E-04 |
| 8BL | 9.60E-03 | 1.08E+01 | 1.79E-25 | 1.33E-01 | 3.36E-02 | 8.87E-03 | 6.44E+00 | 4.07E-10 | 2.38E-01 | 8.94E-04 |
| 9p | 2.17E-02 | 2.13E+01 | 2.62E-82 | 1.24E-01 | 4.64E-02 | 2.70E-02 | 1.85E+01 | 1.40E-64 | 2.08E-01 | 4.70E-03 |
| 10d | 5.75E-03 | 5.82E+00 | 1.42E-08 | 1.72E-01 | 1.45E-02 | 9.38E-03 | 6.48E+00 | 3.11E-10 | 1.48E-01 | 2.37E-02 |
| 8C | -7.89E-03 | -6.70E+00 | 6.98E-11 | 7.27E-02 | 1.86E-01 | -3.56E-03 | -2.37E+00 | 2.17E-02 | 9.57E-02 | 1.12E-01 |
| 44 | 2.27E-02 | 1.56E+01 | 1.42E-48 | 1.40E-01 | 2.58E-02 | 2.93E-03 | 1.63E+00 | 1.14E-01 | 1.61E-01 | 1.81E-02 |
| 45 | 5.12E-02 | 4.01E+01 | 1.15E-209 | 8.46E-02 | 1.40E-01 | 1.91E-02 | 1.14E+01 | 2.09E-27 | 9.24E-02 | 1.08E-01 |
| 47l | 5.12E-03 | 4.61E+00 | 7.32E-06 | 4.75E-02 | 2.87E-01 | -1.10E-02 | -6.65E+00 | 1.11E-10 | 1.66E-01 | 2.19E-02 |
| a47r | -5.25E-03 | -5.10E+00 | 6.62E-07 | 2.04E-01 | 7.50E-03 | -3.19E-03 | -2.37E+00 | 2.17E-02 | 1.31E-02 | 5.06E-01 |
| 6r | -1.25E-02 | -1.15E+01 | 3.73E-28 | 1.53E-01 | 2.24E-02 | -2.59E-02 | -1.86E+01 | 3.30E-65 | 1.28E-01 | 4.93E-02 |
| IFJa | 1.42E-02 | 7.39E+00 | 7.03E-13 | 8.46E-02 | 1.40E-01 | 9.91E-04 | 4.85E-01 | 6.46E-01 | 3.06E-02 | 4.08E-01 |
| IFJp | -3.99E-04 | -2.61E-01 | 8.31E-01 | 1.98E-01 | 3.61E-03 | -3.23E-03 | -1.99E+00 | 5.44E-02 | 1.14E-01 | 4.95E-02 |
| IFSp | 1.77E-02 | 9.94E+00 | 8.86E-22 | 8.95E-02 | 1.28E-01 | 2.75E-04 | 1.41E-01 | 8.93E-01 | 1.57E-01 | 2.61E-02 |
| IFSa | 8.37E-03 | 6.68E+00 | 8.04E-11 | 2.07E-01 | 5.28E-03 | -1.02E-02 | -7.16E+00 | 4.16E-12 | 9.18E-02 | 1.27E-01 |
| p9-46v | -3.91E-02 | -3.45E+01 | 3.85E-171 | 1.73E-01 | 1.50E-02 | -2.86E-02 | -2.18E+01 | 7.44E-85 | 1.60E-01 | 1.59E-02 |
| 46 | -3.03E-02 | -2.81E+01 | 1.52E-127 | 1.88E-01 | 1.03E-02 | -3.11E-02 | -2.30E+01 | 6.23E-93 | 1.46E-01 | 2.59E-02 |
| a9-46v | -2.03E-02 | -1.66E+01 | 3.49E-54 | 1.37E-01 | 2.66E-02 | -1.14E-02 | -8.06E+00 | 6.74E-15 | 1.11E-01 | 5.66E-02 |
| 9-46d | -3.16E-02 | -2.49E+01 | 8.92E-106 | 1.77E-01 | 9.95E-03 | -2.93E-02 | -2.28E+01 | 8.22E-92 | 1.51E-01 | 2.23E-02 |
| 9a | 1.52E-02 | 1.42E+01 | 4.09E-41 | 5.32E-02 | 2.57E-01 | 9.16E-03 | 5.84E+00 | 1.29E-08 | 2.21E-01 | 2.91E-03 |
| 10v | 2.63E-02 | 2.21E+01 | 8.50E-88 | 6.10E-02 | 2.14E-01 | 8.75E-03 | 5.94E+00 | 7.70E-09 | 1.22E-01 | 4.95E-02 |
| a10p | 1.05E-02 | 6.79E+00 | 3.85E-11 | 2.02E-01 | 2.75E-03 | 1.60E-02 | 8.49E+00 | 2.67E-16 | 1.86E-01 | 8.28E-03 |
| 10pp | 4.55E-03 | 3.17E+00 | 2.19E-03 | 1.15E-01 | 6.52E-02 | -3.51E-04 | -1.86E-01 | 8.67E-01 | 1.63E-01 | 1.92E-02 |
| 11l | -5.91E-02 | -4.07E+01 | 6.02E-213 | 3.08E-01 | 1.80E-05 | -4.34E-02 | -2.68E+01 | 7.60E-118 | 2.62E-01 | 1.98E-04 |
| 13l | 2.26E-02 | 1.35E+01 | 1.39E-37 | 1.47E-01 | 2.76E-02 | 4.34E-03 | 2.29E+00 | 2.71E-02 | 1.85E-01 | 1.42E-02 |
| OFC | -7.27E-04 | -4.28E-01 | 7.17E-01 | 1.07E-01 | 7.54E-02 | 2.06E-03 | 9.70E-01 | 3.56E-01 | 4.13E-02 | 3.48E-01 |
| 47s | 2.77E-02 | 2.26E+01 | 4.59E-91 | 1.80E-01 | 1.45E-02 | 1.67E-02 | 1.09E+01 | 1.36E-25 | 1.54E-01 | 2.32E-02 |
| LIPd | -9.51E-03 | -7.72E+00 | 6.73E-14 | 6.44E-02 | 2.06E-01 | -6.57E-03 | -4.36E+00 | 2.12E-05 | 8.00E-02 | 1.40E-01 |
| 6a | -8.19E-03 | -8.42E+00 | 3.31E-16 | 1.76E-01 | 1.07E-02 | -1.89E-02 | -1.44E+01 | 6.61E-42 | 1.53E-01 | 1.86E-02 |
| i6-8 | -1.17E-02 | -1.06E+01 | 1.75E-24 | 7.95E-02 | 1.45E-01 | 1.69E-03 | 1.25E+00 | 2.28E-01 | 1.45E-01 | 3.15E-02 |
| s6-8 | -2.00E-02 | -1.74E+01 | 1.10E-58 | 1.80E-01 | 1.40E-02 | -9.76E-03 | -6.60E+00 | 1.45E-10 | 1.13E-01 | 7.48E-02 |
| 43 | -7.11E-03 | -7.00E+00 | 9.97E-12 | 6.34E-02 | 2.02E-01 | -9.25E-03 | -5.21E+00 | 3.94E-07 | 2.38E-02 | 4.36E-01 |
| OP4 | -3.79E-03 | -3.89E+00 | 1.55E-04 | 1.20E-01 | 6.53E-02 | -6.92E-03 | -3.69E+00 | 3.32E-04 | 0.00E+00 | 5.42E-01 |
| OP1 | -5.49E-03 | -5.59E+00 | 5.24E-08 | 1.73E-01 | 1.62E-02 | -8.83E-03 | -4.74E+00 | 4.04E-06 | 5.30E-02 | 2.97E-01 |
| OP2-3 | -5.61E-03 | -6.13E+00 | 2.42E-09 | 1.63E-01 | 2.44E-02 | -4.41E-03 | -2.67E+00 | 9.54E-03 | 2.77E-02 | 4.19E-01 |
| 52 | -9.24E-03 | -9.71E+00 | 6.70E-21 | 8.74E-02 | 1.23E-01 | -1.02E-02 | -6.16E+00 | 2.13E-09 | 8.02E-02 | 1.65E-01 |
| RI | -1.20E-03 | -1.32E+00 | 2.17E-01 | 1.45E-01 | 2.59E-02 | -1.58E-03 | -9.04E-01 | 3.88E-01 | 9.69E-02 | 1.09E-01 |
| PFcm | -7.49E-03 | -7.19E+00 | 2.71E-12 | 3.88E-02 | 3.30E-01 | -1.13E-02 | -6.24E+00 | 1.35E-09 | 0.00E+00 | 5.42E-01 |
| Pol2 | -5.33E-03 | -5.13E+00 | 5.92E-07 | 8.65E-02 | 1.39E-01 | -8.16E-03 | -4.61E+00 | 7.15E-06 | 0.00E+00 | 5.42E-01 |
| TA2 | -3.24E-03 | -3.01E+00 | 3.73E-03 | 1.41E-01 | 2.75E-02 | -3.64E-03 | -2.24E+00 | 3.06E-02 | 2.35E-02 | 4.34E-01 |
| FOP4 | -1.58E-02 | -1.58E+01 | 1.59E-49 | 4.87E-02 | 2.63E-01 | -1.98E-02 | -1.43E+01 | 3.98E-41 | 1.03E-01 | 1.08E-01 |
| MI | -1.28E-02 | -1.23E+01 | 9.92E-32 | 4.10E-02 | 3.19E-01 | -1.51E-02 | -1.05E+01 | 8.58E-24 | 3.54E-02 | 3.95E-01 |
| Pir | -2.43E-04 | -1.75E-01 | 8.86E-01 | 8.38E-02 | 1.34E-01 | 5.36E-03 | 2.96E+00 | 4.09E-03 | 6.17E-02 | 2.32E-01 |
| AVI | -2.27E-02 | -1.73E+01 | 6.01E-58 | 3.74E-02 | 3.27E-01 | -1.75E-02 | -1.25E+01 | 1.86E-32 | 1.67E-01 | 1.44E-02 |
| AAIC | -1.09E-02 | -6.92E+00 | 1.68E-11 | 2.15E-02 | 4.32E-01 | 5.50E-03 | 2.92E+00 | 4.68E-03 | 6.64E-02 | 2.17E-01 |
| FOP1 | -1.45E-02 | -1.28E+01 | 5.44E-34 | 7.44E-02 | 1.87E-01 | -1.72E-02 | -1.02E+01 | 8.88E-23 | 1.00E-01 | 1.24E-01 |
| FOP3 | -1.07E-02 | -1.05E+01 | 3.10E-24 | 0.00E+00 | 5.26E-01 | -1.45E-02 | -9.36E+00 | 2.17E-19 | 9.86E-03 | 5.28E-01 |
| FOP2 | -3.75E-03 | -3.80E+00 | 2.19E-04 | 9.68E-02 | 1.12E-01 | -1.05E-02 | -5.96E+00 | 6.66E-09 | 0.00E+00 | 5.42E-01 |
| PFt | 2.59E-03 | 2.94E+00 | 4.50E-03 | 1.62E-01 | 2.19E-02 | -6.70E-03 | -4.68E+00 | 5.19E-06 | 5.42E-02 | 2.80E-01 |
| AIP | -6.30E-04 | -7.22E-01 | 5.16E-01 | 1.28E-01 | 4.38E-02 | -6.90E-03 | -5.45E+00 | 1.14E-07 | 1.57E-02 | 4.79E-01 |
| EC | 4.12E-03 | 2.21E+00 | 3.48E-02 | 5.42E-02 | 2.46E-01 | 5.79E-03 | 2.70E+00 | 8.77E-03 | 3.83E-02 | 3.57E-01 |
| PreS | 1.23E-02 | 7.96E+00 | 1.19E-14 | 8.15E-02 | 1.29E-01 | 3.53E-02 | 1.89E+01 | 2.42E-67 | 3.03E-02 | 4.08E-01 |
| H | 8.89E-03 | 5.19E+00 | 4.23E-07 | 7.57E-02 | 1.64E-01 | 6.36E-03 | 3.42E+00 | 8.98E-04 | 2.65E-02 | 4.15E-01 |
| ProS | 7.61E-03 | 6.75E+00 | 4.89E-11 | 6.54E-02 | 2.05E-01 | 2.05E-02 | 1.35E+01 | 3.36E-37 | 7.01E-02 | 2.21E-01 |
| PeEc | -1.89E-03 | -1.24E+00 | 2.47E-01 | 7.97E-02 | 1.63E-01 | -7.96E-03 | -4.45E+00 | 1.46E-05 | 7.54E-02 | 1.78E-01 |
| STGa | 3.75E-02 | 2.32E+01 | 7.81E-95 | 2.91E-02 | 3.90E-01 | -9.15E-03 | -5.04E+00 | 9.10E-07 | 0.00E+00 | 5.42E-01 |
| PBelt | -5.69E-03 | -5.88E+00 | 1.03E-08 | 2.01E-01 | 5.66E-03 | -5.13E-03 | -3.10E+00 | 2.65E-03 | 8.70E-02 | 1.47E-01 |
| A5 | 2.24E-02 | 1.61E+01 | 1.76E-51 | 1.97E-01 | 4.23E-03 | -2.64E-03 | -1.55E+00 | 1.32E-01 | 3.55E-02 | 3.88E-01 |
| PHA1 | 4.02E-03 | 2.65E+00 | 1.07E-02 | 9.45E-02 | 1.03E-01 | 1.98E-02 | 1.10E+01 | 4.25E-26 | 1.01E-01 | 1.08E-01 |
| PHA3 | 1.87E-03 | 1.61E+00 | 1.28E-01 | 9.75E-02 | 9.77E-02 | 6.15E-03 | 3.90E+00 | 1.48E-04 | 2.68E-02 | 4.18E-01 |
| STSda | 1.70E-02 | 1.12E+01 | 6.97E-27 | 1.01E-01 | 8.66E-02 | -1.28E-02 | -7.01E+00 | 1.10E-11 | 5.85E-02 | 2.37E-01 |
| STSdp | 5.53E-02 | 3.41E+01 | 7.24E-169 | 4.91E-02 | 2.62E-01 | 1.04E-02 | 5.51E+00 | 8.24E-08 | 1.51E-01 | 2.42E-02 |
| STSvp | 1.06E-02 | 1.12E+01 | 7.76E-27 | 1.54E-01 | 2.17E-02 | 5.36E-03 | 3.86E+00 | 1.72E-04 | 2.02E-01 | 5.37E-03 |
| TGd | 2.81E-02 | 2.30E+01 | 1.85E-93 | 5.56E-02 | 2.43E-01 | 9.17E-03 | 5.87E+00 | 1.10E-08 | 1.58E-01 | 1.88E-02 |
| TE1a | 1.78E-02 | 1.86E+01 | 1.10E-65 | 5.76E-02 | 2.31E-01 | 1.62E-02 | 1.19E+01 | 9.45E-30 | 1.97E-01 | 3.89E-03 |
| TE1p | -6.01E-03 | -5.92E+00 | 8.22E-09 | 1.46E-01 | 2.22E-02 | 2.22E-03 | 1.74E+00 | 9.22E-02 | 1.22E-01 | 4.88E-02 |
| TE2a | -5.75E-03 | -5.40E+00 | 1.45E-07 | 1.17E-01 | 6.71E-02 | -1.07E-02 | -7.21E+00 | 3.03E-12 | 6.60E-02 | 2.11E-01 |
| TF | 2.19E-02 | 1.28E+01 | 2.35E-34 | 1.94E-01 | 3.54E-03 | 1.17E-02 | 5.86E+00 | 1.16E-08 | 1.37E-01 | 2.79E-02 |
| TE2p | 1.30E-02 | 9.53E+00 | 3.44E-20 | 2.11E-01 | 3.27E-03 | 2.35E-03 | 1.47E+00 | 1.53E-01 | 1.41E-01 | 2.94E-02 |
| PHT | -6.93E-03 | -6.81E+00 | 3.43E-11 | 1.23E-01 | 4.68E-02 | -2.01E-02 | -1.47E+01 | 3.44E-43 | 1.19E-01 | 5.57E-02 |
| PH | 9.73E-03 | 8.98E+00 | 3.62E-18 | 5.44E-02 | 2.62E-01 | 1.35E-02 | 8.06E+00 | 6.84E-15 | 0.00E+00 | 5.42E-01 |
| TPOJ1 | 1.01E-02 | 5.77E+00 | 1.86E-08 | 1.90E-01 | 8.17E-03 | 1.74E-03 | 9.55E-01 | 3.62E-01 | 2.10E-01 | 4.23E-03 |
| TPOJ2 | -3.58E-03 | -2.71E+00 | 8.98E-03 | 2.14E-01 | 1.76E-03 | -1.22E-03 | -7.71E-01 | 4.59E-01 | 2.18E-01 | 2.12E-03 |
| TPOJ3 | -3.02E-03 | -2.07E+00 | 4.82E-02 | 1.84E-02 | 4.42E-01 | 3.23E-03 | 1.94E+00 | 6.10E-02 | 9.82E-02 | 1.07E-01 |
| DVT | -3.73E-03 | -4.12E+00 | 6.24E-05 | 9.77E-02 | 1.08E-01 | 5.66E-03 | 4.37E+00 | 2.08E-05 | 3.53E-03 | 5.71E-01 |
| PGp | -8.30E-03 | -7.86E+00 | 2.37E-14 | 1.39E-01 | 2.62E-02 | -7.66E-03 | -5.46E+00 | 1.08E-07 | 8.69E-02 | 1.27E-01 |
| IP2 | -2.52E-02 | -2.46E+01 | 3.76E-104 | 1.62E-01 | 1.51E-02 | -1.31E-02 | -1.08E+01 | 6.88E-25 | 1.75E-01 | 7.96E-03 |
| IP1 | -2.87E-02 | -2.97E+01 | 1.79E-138 | 1.07E-01 | 8.36E-02 | -2.06E-02 | -1.73E+01 | 1.15E-57 | 1.54E-01 | 1.88E-02 |
| IP0 | 1.75E-03 | 1.53E+00 | 1.49E-01 | 6.09E-02 | 2.28E-01 | 3.04E-03 | 1.85E+00 | 7.29E-02 | 0.00E+00 | 5.42E-01 |
| PFop | -2.25E-03 | -2.07E+00 | 4.76E-02 | 1.20E-02 | 4.87E-01 | -8.94E-03 | -5.19E+00 | 4.33E-07 | 0.00E+00 | 5.42E-01 |
| PF | -8.73E-03 | -8.84E+00 | 1.15E-17 | 1.41E-01 | 2.90E-02 | -2.01E-02 | -1.44E+01 | 5.68E-42 | 9.44E-02 | 1.26E-01 |
| PFm | -3.51E-03 | -4.07E+00 | 7.54E-05 | 8.95E-02 | 1.27E-01 | 1.29E-04 | 1.08E-01 | 9.14E-01 | 5.51E-02 | 2.77E-01 |
| PGi | 1.39E-02 | 1.54E+01 | 2.62E-47 | 1.12E-01 | 6.47E-02 | 1.14E-02 | 8.36E+00 | 7.20E-16 | 1.98E-01 | 5.15E-03 |
| PGs | -7.72E-05 | -1.03E-01 | 9.28E-01 | 9.39E-02 | 1.21E-01 | 1.30E-02 | 1.06E+01 | 3.93E-24 | 1.48E-01 | 2.57E-02 |
| V6A | -3.71E-04 | -3.55E-01 | 7.65E-01 | 2.99E-02 | 3.87E-01 | 1.19E-02 | 6.98E+00 | 1.30E-11 | 0.00E+00 | 5.42E-01 |
| VMV<br>1 | 2.32E-03 | 2.39E+00 | 2.19E-02 | 0.00E+00 | 5.26E-01 | 1.37E-02 | 8.87E+00 | 1.21E-17 | 0.00E+00 | 5.42E-01 |
| VMV<br>3 | 1.05E-03 | 9.26E-01 | 3.97E-01 | 0.00E+00 | 5.26E-01 | 1.14E-02 | 6.06E+00 | 3.92E-09 | 0.00E+00 | 5.42E-01 |
| PHA2 | 1.93E-02 | 1.34E+01 | 2.90E-37 | 1.62E-01 | 1.45E-02 | 3.18E-02 | 1.81E+01 | 3.26E-62 | 1.48E-01 | 2.90E-02 |
| V4t | -2.62E-03 | -2.44E+00 | 1.93E-02 | 0.00E+00 | 5.26E-01 | 1.20E-02 | 7.01E+00 | 1.11E-11 | 3.25E-02 | 4.06E-01 |
| FST | 2.62E-03 | 2.74E+00 | 8.37E-03 | 8.76E-02 | 1.29E-01 | 1.12E-02 | 7.43E+00 | 6.75E-13 | 2.20E-02 | 4.45E-01 |
| V3CD | 1.47E-03 | 1.30E+00 | 2.24E-01 | 0.00E+00 | 5.26E-01 | 1.34E-02 | 6.78E+00 | 5.05E-11 | 1.96E-02 | 4.66E-01 |
| LO3 | 1.67E-04 | 1.52E-01 | 8.99E-01 | 0.00E+00 | 5.26E-01 | 1.22E-02 | 6.77E+00 | 5.12E-11 | 0.00E+00 | 5.42E-01 |
| VMV<br>2 | 4.62E-03 | 4.40E+00 | 1.84E-05 | 0.00E+00 | 5.26E-01 | 1.28E-02 | 7.52E+00 | 3.61E-13 | 0.00E+00 | 5.42E-01 |
| 31pd | 1.29E-02 | 1.26E+01 | 4.36E-33 | 7.52E-02 | 1.64E-01 | 1.82E-02 | 1.20E+01 | 2.15E-30 | 8.24E-02 | 1.32E-01 |
| 31a | -1.65E-02 | -1.35E+01 | 1.18E-37 | 5.74E-02 | 2.30E-01 | 2.56E-03 | 1.75E+00 | 9.00E-02 | 0.00E+00 | 5.42E-01 |
| VVC | 7.73E-04 | 6.86E-01 | 5.38E-01 | 0.00E+00 | 5.26E-01 | 1.04E-02 | 5.62E+00 | 4.63E-08 | 1.95E-03 | 5.80E-01 |
| 25 | 1.49E-03 | 9.44E-01 | 3.91E-01 | 8.29E-02 | 1.40E-01 | 1.10E-03 | 6.08E-01 | 5.62E-01 | 2.74E-02 | 4.16E-01 |
| s32 | 1.08E-03 | 8.23E-01 | 4.54E-01 | 6.73E-02 | 1.78E-01 | -1.35E-03 | -7.86E-01 | 4.55E-01 | 1.36E-01 | 3.36E-02 |
| pOFC | -3.37E-03 | -1.71E+00 | 1.06E-01 | 6.96E-02 | 1.84E-01 | 5.77E-03 | 2.66E+00 | 9.79E-03 | 4.99E-02 | 3.11E-01 |
| PoI1 | -7.53E-03 | -7.03E+00 | 8.26E-12 | 2.72E-02 | 4.05E-01 | -1.20E-02 | -7.35E+00 | 1.16E-12 | 7.77E-02 | 1.88E-01 |
| Ig | -9.35E-05 | -9.78E-02 | 9.27E-01 | 1.29E-01 | 5.45E-02 | -3.24E-03 | -1.79E+00 | 8.40E-02 | 7.81E-02 | 1.78E-01 |
| FOP5 | -2.37E-02 | -1.73E+01 | 3.01E-58 | 1.36E-01 | 3.10E-02 | -2.56E-02 | -1.68E+01 | 1.13E-54 | 1.85E-01 | 1.27E-02 |
| p10p | -2.81E-02 | -2.22E+01 | 5.14E-88 | 1.97E-01 | 7.73E-03 | -1.75E-02 | -1.08E+01 | 3.79E-25 | 1.45E-01 | 2.66E-02 |
| p47r | -1.59E-02 | -1.40E+01 | 2.90E-40 | 1.80E-01 | 8.17E-03 | -1.14E-02 | -8.36E+00 | 7.17E-16 | 1.14E-01 | 6.29E-02 |
| TGv | 5.67E-02 | 3.40E+01 | 2.33E-168 | 2.48E-02 | 4.09E-01 | 1.84E-02 | 9.28E+00 | 3.92E-19 | 2.82E-02 | 4.07E-01 |
| MBelt | -1.12E-02 | -1.11E+01 | 9.38E-27 | 2.44E-01 | 9.36E-04 | -9.80E-03 | -6.08E+00 | 3.37E-09 | 1.06E-01 | 9.18E-02 |
| LBelt | -6.00E-03 | -6.51E+00 | 2.34E-10 | 2.66E-01 | 4.98E-04 | -5.25E-03 | -3.40E+00 | 9.64E-04 | 1.72E-01 | 1.77E-02 |
| A4 | 2.60E-04 | 2.52E-01 | 8.29E-01 | 1.59E-01 | 1.62E-02 | -4.37E-03 | -2.62E+00 | 1.10E-02 | 5.99E-02 | 2.55E-01 |
| STSva | 1.26E-02 | 1.13E+01 | 1.83E-27 | 8.61E-02 | 1.26E-01 | 5.11E-03 | 3.26E+00 | 1.53E-03 | 2.42E-01 | 2.99E-04 |
| TE1m | 2.35E-03 | 2.57E+00 | 1.33E-02 | 1.46E-01 | 2.75E-02 | 6.26E-03 | 5.00E+00 | 1.10E-06 | 7.26E-02 | 1.89E-01 |
| PI | -1.00E-02 | -8.94E+00 | 4.90E-18 | 5.73E-02 | 2.47E-01 | -6.97E-03 | -4.38E+00 | 2.02E-05 | 3.21E-02 | 4.09E-01 |
| a32pr | -1.17E-02 | -9.16E+00 | 7.60E-19 | 1.51E-01 | 2.26E-02 | -1.65E-02 | -1.21E+01 | 6.39E-31 | 1.56E-01 | 2.29E-02 |
| p24 | -1.06E-02 | -8.11E+00 | 3.67E-15 | 3.41E-02 | 3.72E-01 | -6.44E-03 | -4.59E+00 | 7.80E-06 | 1.14E-01 | 8.56E-02 |

## Notes

### Competing Interest Statement

The authors have declared no competing interest.

